# CK1 Delta Is an mRNA Cap-Associated Protein That Drives Translation Initiation and Tumor Growth

**DOI:** 10.1101/2020.02.20.955229

**Authors:** Ipsita Pal, Andre M. Sardinha Grilo, Luke E. Berchowitz, Sohani Das Sharma, Prabhjot S. Mundi, Abdullah M. Ali, M. Laura Martin, Maryam Safari, Luigi Scotto, Eli Malkovskiy, Ahmed Sawas, Siddhartha Mukherjee, Owen A. O’Connor, Marko Jovanovic, Changchun Deng

**Affiliations:** Center for Lymphoid Malignancies, Department of Medicine, Columbia University Irving Medical Center; Department of Genetics and Development, Columbia University; Department of Biological Sciences, Columbia University; Division of Hematology and Oncology, Department of Medicine, Columbia University Irving Medical Center; Institute of Cancer Genetics, Columbia University Irving Medical Center; Physiology and Biophysics, Englander Institute for Precision Medicine, Weill Cornell Medical College

## Abstract

Whether translation is differentially regulated across liquid and solid tumors remains poorly understood. Here we report the discovery that Casein Kinase 1 delta (CK1δ) plays a key role in regulating translation initiation in blood cancers, but interestingly, not in solid tumors. In lymphomas CK1δ is a key positive regulator of 4E-BP1 and p70S6K phosphorylation, assembly of eIF4F, and translation initiation. Furthermore, CK1δ is pulled down by m^7^GTP-agarose that mimics the mRNA m^7^G cap, consistent with the regulatory role of CK1δ in translation initiation. Targeting CK1δ using a small molecule inhibitor, namely SR-3029, potently kills lymphoma cell lines and primary lymphoma cells across histology subtypes. While SR-3029 shares with mTORC1 inhibitors the overlapping mechanism of repressing 4E-BP1 and p70S6K/RPS6 phosphorylation, the kinetics of repression is slow with SR-3029 and fast with mTORC1 inhibitors such as Torin-1. Remarkably, it is slower-acting SR-3029, but not fast-acting Torin-1, that kills lymphoma cells consistently across multiple histology subtypes. Proteomics and RNA sequencing studies show that SR-3029 represses the expression of many genes preferentially at the translation step, such as genes in the reactome translation initiation pathway. SR-3029 markedly represses the protein level of the C-MYC oncogene without decreasing its mRNA level. In contrast, Torin-1 fails to reduce the protein level of C-MYC in the same lymphoma cells. While SR-3029 also demonstrates potent activity in select solid tumors, its mechanism of action in the solid tumors is different. In breast cancer cells SR-3029 inhibits nuclear localization of β-catenin but at the same concentrations does not inhibit 4E-BP1 and p70S6K phosphorylation or global protein synthesis. Likewise, SR-3029 does not inhibit nuclear localization of β-catenin in lymphoma cells. Our results indicate that CK1δ is an mRNA cap-associated protein and an upstream kinase required for 4E-BP1 and p70S6K phosphorylation. CK1δ stimulates assembly of eIF4F and translation initiation, and is a critical driver for tumor growth in blood cancers across multiple histology types. CK1δ invokes the alternative mechanism of regulating β-catenin in select solid tumors. Our results indicate that CK1δ inhibition is a promising therapeutic strategy in both liquid and solid tumors, and the distinct roles of CK1δ in these malignancies may serve as biomarkers to enable precision cancer treatment.

## INTRODUCTION

Global protein synthesis rate plays a critical role in the governance of cell growth, cell size, and tumor cell proliferation. Targeting translation, especially initiation, could thus be an effective strategy to target and treat fast-growing cancers. Stabilization of the eukaryotic translation initiation factor 4F (eIF4F) complex is a hallmark of resistance to anti-BRAF and anti-MEK therapies in several melanoma and colon cancer models [1]. This often occurs through persistent hyperphosphorylation and thus inactivation of 4E-BP1, a negative regulator of eIF4F. Our recent studies have strongly implicated that targeting of 4E-BP1 is a viable strategy for the treatment of several aggressive lymphomas [2]. Here, we identify Casein Kinase 1 delta (CK1δ) as an mRNA cap-associated protein, a critical regulator of 4E-BP1 phosphorylation, and activator of mRNA translation. Furthermore, we demonstrate that chemical inhibition of this kinase potently inhibits tumor growth in models of blood cancers.

We recently showed that combination therapy of TGR-1202 and carfilzomib synergistically inhibits phosphorylation of 4E-BP1, translation of the C-MYC oncogene, and cell viability in lymphoma and myeloma cells [2]. This regimen is now being studied in a phase I clinical trial (NCT02867618). Interestingly, this regimen depends on TGR-1202 to inhibit PI3Kδ, its target by design, and CK1 epsilon (CK1ε), an unexpected target [2]. CK1ε is reported to regulate translation initiation via phosphorylating 4E-BP1 [3]. Surprisingly, when we knocked out or chemically inhibited CK1ε in lymphoma cells we did not observe any change in the phosphorylation of 4E-BP1. These seemingly contradictory results may be reconciled at least in part by the fact that a potent dual CK1ε/CK1δ inhibitor (PF670462), but not the CK1ε inhibitor PF4800567, was utilized to establish the role of CK1ε in regulating phosphorylation of 4E-BP1 and translation in the previous study [3]. The discrepancy between our results [2] and the previous report [3] thus led us to investigate here the hypothesis that CK1δ is a key regulator of 4E-BP1 and translation.

The role of CK1δ in translation is poorly defined, although it has been reported that CK1ε and CK1δ collectively are required for pre-40S ribosome maturation [4]. CK1δ has been reported to mediate Wnt/β-catenin signaling in breast cancer, and selective inhibition of CK1δ using the molecule SR-3029 [5] potently kills breast cancer cell lines that exhibit high CK1δ protein levels via downregulating Wnt/β-catenin signaling[6].

In this study we demonstrate that CK1δ regulates translation by mediating phosphorylation of 4E-BP1 and p70S6K. We show that CK1δ localizes to the 7-methylguanylate (m^7^G) structure at the 5’ end of mRNA cap. Furthermore, chemical inhibition of CK1δ using SR-3029 potently downregulates 4E-BP1 and p70S6K/RPS6 phosphorylation and disrupts eIF4F assembly. Therapeutically SR-3029 potently reduces blood cancer cell viability in a wide array of models. SR-3029 is often superior to the mTORC1/2 inhibitor Torin-1, which targets translation initiation via inhibiting phosphorylation of 4E-BP1 and p70S6K. The CK1δ inhibitor SR-3029 is very effective in breast cancer and colon cancer models, but not in pancreatic cancer. In breast cancer cells SR-3029 does not inhibit phosphorylation of 4E-BP1 and p70S6K/RPS6, but does inhibit nuclear localization of β-catenin, which is in agreement with previous results [6]. Likewise, in lymphoma cells SR-3029 does not inhibit nuclear localization of β-catenin. These results highlight that the anti-tumor activity of SR-3029 acts via differential pathways in liquid vs. solid tumors. Our results demonstrate that CK1δ is a key regulator of translation and is a promising therapeutic target in lymphomas.

## RESULTS

### CK1δ is required for phosphorylation of 4E-BP1, assembly of eIF4F, and efficient translation

Our first goal was to test the hypothesis that CK1δ regulates 4E-BP1 phosphorylation and activates mRNA translation. We used CRISPR/Cas9 to generate four single cell-derived clones of CK1δ knockout (CK1δ-/-) in HEK293 cells, and the clones were sequenced to confirm the target specificity. Our four clones KO1-4 had no detectable CK1δ protein but showed normal levels of CK1ε (**Figure 1A**). We investigated whether the absence of CK1δ affected phosphorylation of 4E-BP1. Phosphorylation of 4E-BP1 was markedly decreased in the CK1δ-/- clones KO3 and KO4 but not affected in KO1 and KO2 (**Figure 1A**). The protein level of total 4E-BP1 in the CK1δ-/- clones KO3 and KO4 was mildly reduced compared to the control HEK293 cells containing only Cas9. In contrast, when CK1ε was knocked out, we observed no significant change in the protein levels of 4E-BP1 and phosphorylated 4E-BP1 (**Figure S1A**). These results demonstrate that CK1δ regulates phosphorylation of 4E-BP1 in HEK293. Similar to 4E-BP1 the kinase p70S6K is phosphorylated by mTOR (mechanistic target of rapamycin) during translation initiation [7-10]. We reasoned that loss of CK1δ may inhibit phosphorylation of p70S6K. Surprisingly, phosphorylation of p70S6K or its substrate RPS6 was not affected in the CK1δ-/- clones (**Figure 1B**). These results demonstrate that CK1δ is required specifically for phosphorylation of 4E-BP1, but does not regulate p70S6K/RPS6 phosphorylation in HEK293.

**Figure 1.**
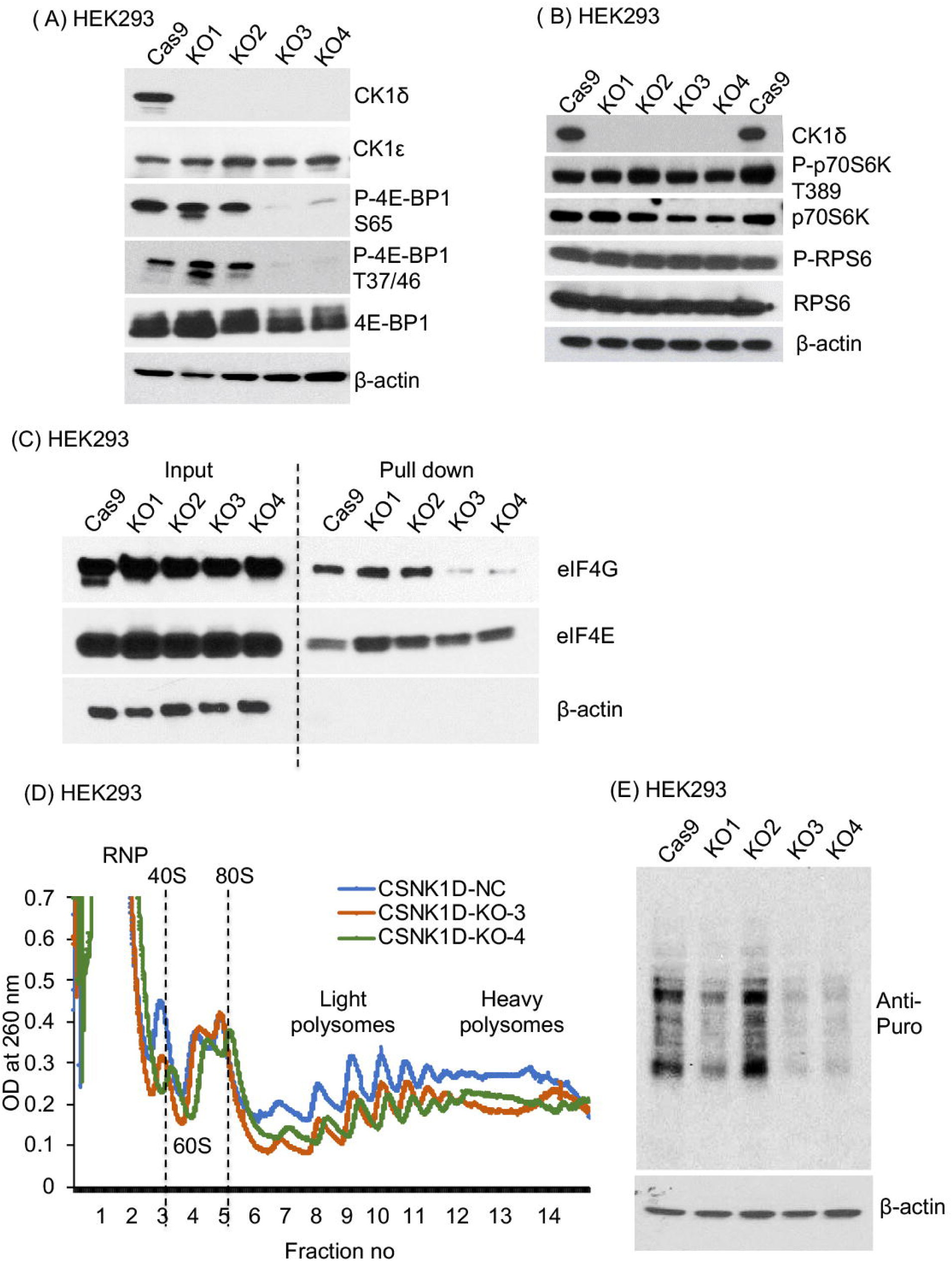
CK1δ is required for phosphorylation of 4E-BP1, assembly of eIF4F, and efficient translation in HEK293. The HEK293 cell line was targeted for CK1δ-/- knockout by the CRISPR-Cas9 system, using 2 different single guide RNA (sgRNA) sequences. Knockout clones were selected from single cells by GFP sorting followed by serial dilution, and gene editing was confirmed by sequencing. KO1 and KO2 had mono-allelic knockout of CK1δ, and KO3 and KO4 has bi-allelic knockout of CK1δ (Figure S1A). **(A-B)** Four clones of CK1δ knockout, KO1-4, were compared with the vector Cas9 in HEK293 for the levels of CK1δ, CK1ε, 4E-BP1, p70S6K, RPS6, and actin by immunoblotting. **(C)** The same CK1δ knockout clones of HEK293 and Cas9 HEK293 cells were processed for the cap binding assay using m^7^GTP agarose beads to pull down cap binding proteins, which were then analyzed by immunoblotting. The ratio of eIF4G in the pulldown relative to the input cell lysate is an index of eIF4F assembly. **(D)** Polysome profiling was performed the in HEK293 cells with CK1δ knockout clones KO3 and KO4, in comparison to the control HEK293 cells with non-targeting Cas9. X-axis showed the sedimentations or fractions collected from 10–50% sucrose density gradient centrifugation. Y-axis showed the OD260 UV recording of RNA abundance. Data shown were the average of three experiments. RNP, ribonuclear protein; 40S, 40S small ribosome subunit; 60S, 60S large ribosome subunit; 80S, 80S monosome. **(E)** The same CK1δ knockout clones of HEK293 and Cas9 HEK293 cells were grown in 1μg/ml puromycin for 30 minutes. Cell lysates were subjected to Western blotting using the anti-puromycin antibody in the SUnSET assay.

We next investigated whether CK1δ knockout directly represses translation, using three independent assays. First, we studied assembly of the eIF4F complex (comprising eIF4G, eIF4E, and eIF4A) using a m^7^GTP cap binding assay[10], in which m^7^GTP-agarose mimics the mRNA m^7^G cap. The level of eIF4G in the pulldown fraction relative to eIF4E, which binds directly to the mRNA m^7^G cap, indicates the efficiency of eIF4F assembly. While the level of eIF4G in the lysate remained constant among the CK1δ-/- clones and Cas9 controls, the level of eIF4G in the pulldown was reduced substantially in KO3 and KO4 (**Figure 1C**). This indicates that CK1δ plays an important role in either assembly or recruitment of eIF4F. We next wanted to test whether CK1δ-/- cells exhibit a defect in global translation. We compared the polysome profiles of CK1δ-/- cells to control cells fractionated in sucrose density gradients. Compared to the control, CK1δ-/- clones KO3 and KO4 exhibited reduced polysome peaks which is indicative of global translation defect (**Figure 1D**). To further analyze global protein synthesis in CK1δ-/- cells we utilized the surface sensing of translation (SUnSET) assay in which puromycin is used as a structural analog of aminoacyl tRNAs[11]. The incorporation of puromycin results in termination of nascent polypeptide chains, which can be monitored using an anti-puromycin antibody. Compared to the Cas9 control, CK1δ-/- clones KO3 and KO4 exhibited reduced puromycin incorporation, or polypeptide synthesis (**Figure 1E**). Collectively the above results support the model that CK1δ activates initiation of translation, likely via mediating phosphorylation of 4E-BP1 thereby promoting assembly of eIF4F.

### CK1δ regulates 4E-BP1/eIF4F in lymphoma and binds to the m^7^G cap-like structure

Given the important role of CK1δ in regulating translation initiation in HEK293 we hypothesized that CK1δ may serve the same function in cancer. We generated a CK1δ knockout (CK1δ-/-) using CRISPR/Cas9 in the human cell line Z-138, which is established from aggressive mantle cell lymphoma (MCL). We utilized mixed populations of three knockout samples for our analysis. The protein level of CK1δ was moderately reduced in the mixed population of KO1 CK1δ-/- cells and near completely reduced in the mixed populations of KO2 and KO3 CK1δ-/- cells. The level of CK1ε in KO1-3 was not altered relative to the Cas9 control (**Figure 2A**). Phosphorylation of 4E-BP1 was mildly reduced in KO1 and markedly reduced in KO2-3 (**Figure 2B**). Furthermore, phosphorylation of RPS6 was substantially reduced in KO3 knockout cells of Z-138 (**Figure 2B**), which is different from the observation in HEK293 knockout clones. We next asked whether chemical inhibition of CK1δ may alter these phosphoproteins. When treated for 24h using SR-3029, a potent and selective inhibitor of CK1δ [5], phosphorylation of 4E-BP1 and p70S6K/RPS6 was markedly reduced in the Z-138 lymphoma cell line (**Figure 2C**). Given the role of 4E-BP1 as a negative regulator of eIF4F, the above results led us to hypothesize that the CK1δ inhibitor SR-3029 will disrupt eIF4F assembly in the Z-138 lymphoma cells. Using the m^7^GTP cap binding assay we observed that SR-3029 treatment substantially reduced eIF4F assembly (**Figure 2D**). Similar results were observed in another MCL cell line Jeko-1 (**Figure S2A**). These above results indicate that CK1δ acts as a positive regulator of 4E-BP1 phosphorylation and eIF4F assembly.

**Figure 2.**
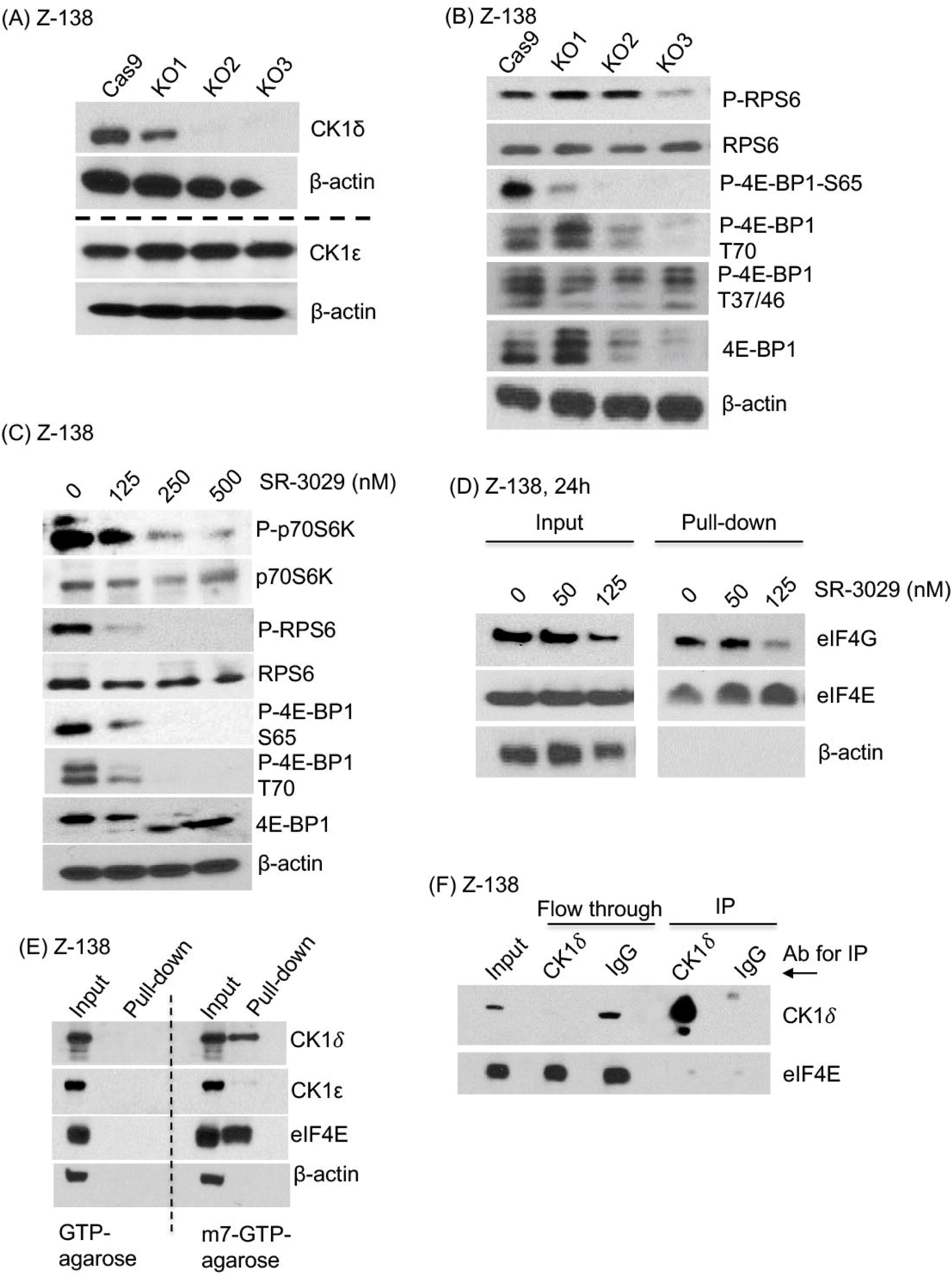
CK1δ regulates 4E-BP1 and eIF4F in lymphoma and binds to the m^7^G cap-like structure. **(A-B)** The MCL cell line Z-138 was targeted for CK1δ knockout by the CRISPR-Cas9 system, using 3 single guide RNA (sgRNA) sequences. Cas9 was the empty vector without any guide sequence. Mixed populations were studied by Western blot using the antibodies as shown in a Western blot. **(C)** Lymphoma cell line Z-138 was treated with SR-3029 or the DMSO control at the indicated concentrations for 24h then processed for immunoblotting. **(D)** Z-138 was treated with SR-3029 or the DMSO control at the indicated concentrations for 24h then processed for the cap binding assay using m^7^GTP agarose beads to pull down cap binding proteins, which were then analyzed by immunoblotting. The ratio of eIF4G in the pulldown relative to the input cell lysate is an index of eIF4F assembly. **(E)** Z-138 cells in logarithmic phase were harvested for lysates, and cap binding assay was performed using m^7^GTP agarose beads and the control GTP-agarose beads. Proteins in the input lysates and the pull-down fractions were analyzed by immunoblot using the antibodies as indicated. **(F)** Immunoprecipitation. Z-138 cell lysates were incubated overnight with anti-CK1δ or a control IgG immobilized to resin, indicated by the arrow. The input, immunoprecipitated sample (IP), and flow through were analyzed by immunoblot using the indicated antibodies.

We asked whether CK1δ is physically associated with eIF4F. Interestingly, we observed that a high proportion of CK1δ was pulled down by the m^7^GTP cap-like structure, which was similarly seen with the cap-binding protein eIF4E (**Figure 2E**). This finding was unexpected, given that a quantitative proteomic study of mRNA cap-associated proteins in HEK293 does not identify CK1δ among 160 plus cap associated proteins [12]. The binding of CK1δ and eIF4E to the m^7^GTP-agarose beads was specific, because the control GTP-agarose beads failed to pull down CK1δ or eIF4E, furthermore, CK1ε was not pulled down by m^7^GTP-agarose (**Figure 2E**). Because CK1δ and eIF4E co-localized in the above cap-binding assay, we asked whether they interact with each other in the cell lysate. Surprisingly, immunoprecipitation using an anti-CK1δ antibody did not precipitate eIF4E (**Figure 2F**). As expected, the anti-CK1δ antibody efficiently and specifically precipitated the designated antigen CK1δ (**Figure 2G**). These results indicate that CK1δ mediates 4E-BP1 and p70S6K/RPS6 phosphorylation, promotes assembly of eIF4F, and is localized or enriched at the mRNA m^7^G cap in lymphoma cells.

### CK1δ inhibitor SR-3029 potently blocks tumor growth and 4E-BP1 phosphorylation

Because the CK1δ inhibitor SR-3029 inhibits eIF4F in the MCL lymphoma cell Z-138, we investigated whether SR-3029 may be a potential treatment for fast growing blood cancers, including MCL and lymphomas of other histology. A recent study reported that the anti-tumor activity of SR-3029 positively correlates with CK1δ protein levels in breast cancer models but it is ineffective in a number of breast cancer cell lines with low CK1δ levels [6]. We found that CK1δ was abundantly expressed in 12 unselected blood cancer cell lines (**Figure S3A**), which led us to hypothesize that SR-3029 may be a useful treatment for these blood cancers regardless of their histology. Using the Cell Titer Glo assay we found that SR-3029 potently inhibited the viability of other human MCL cell lines in addition to Z-138 (**Figure 3A**). Moreover, SR-3029 was highly cytotoxic in other distinct histological types of human lymphomas such as diffuse large B-cell lymphoma (DLBCL) and peripheral T-cell lymphoma (PTCL) (**Figure 3B, 3C**). In contrast, the DLBCL cell line VAL was very resistant to SR-3029 (**Figure 3B**).

**Figure 3.**
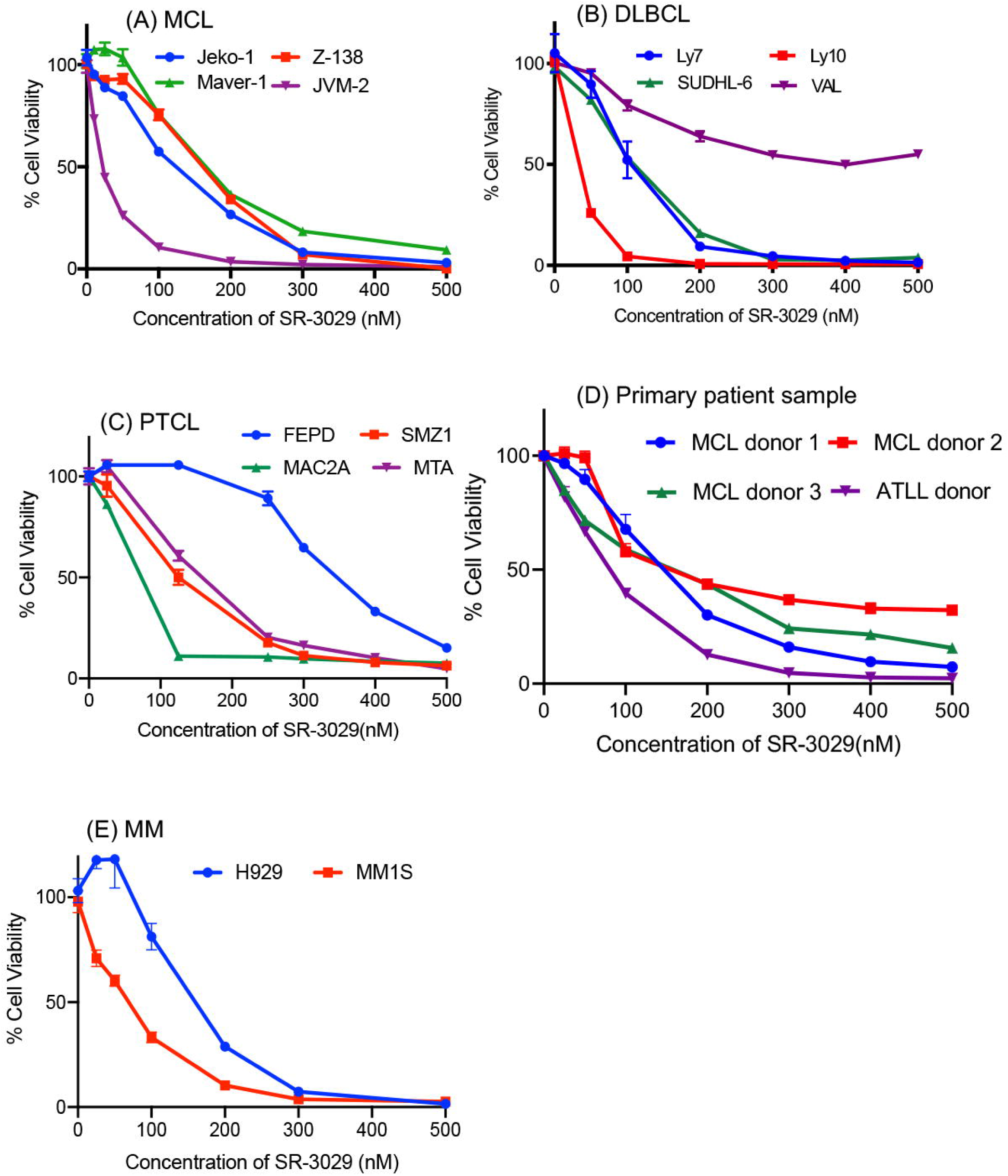
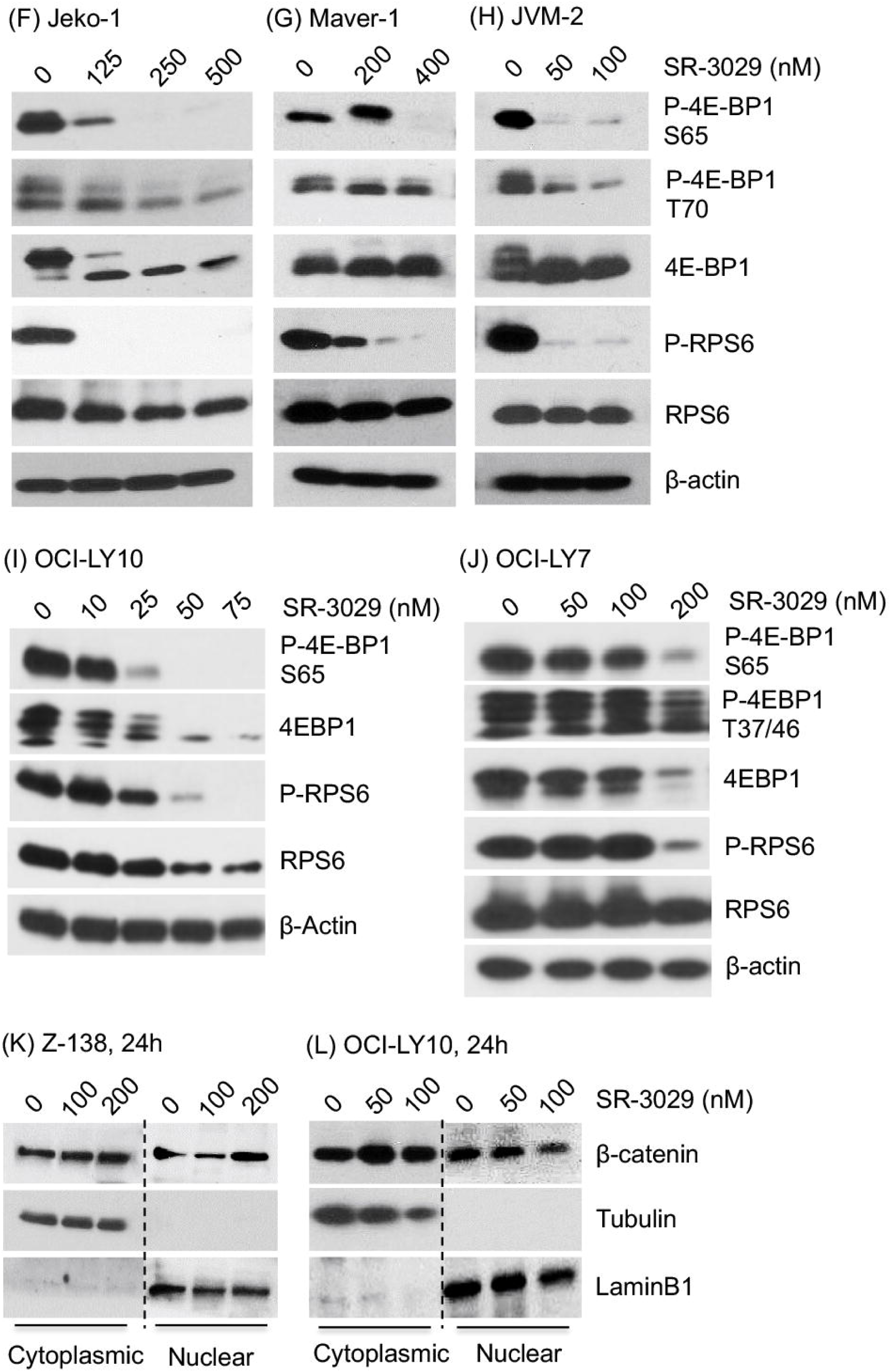
SR-3029 potently inhibits tumor growth in a histology-agnostic manner in blood cancers while repressing phosphorylation of 4E-BP1 and RPS6. **(A-E)** Lymphoma cell lines or primary lymphoma cells were treated with SR-3029 or the DMSO control at the indicated concentrations for 24, 48, and 72h. The viable cells in the treated and control samples were quantitated using the Cell-Titer Glo assay. Only the results from the 48h treatment were shown here. Data shown were the average of three experiments and were presented as mean ± SEM. DLBCL: diffuse large B cell lymphoma, MCL: mantle cell lymphoma, PTCL: peripheral T cell lymphoma, MM: multiple myeloma, ATLL: adult T cell leukemia/lymphoma. **(F-J)** Lymphoma cell lines were treated with SR-3029 for 24h and processed for immunoblotting using the indicated antibodies. **(K-L)** Lymphoma cell lines Z-138 (L) and OCI-LY10 (M) were treated with SR-3029 or DMSO control for 24h as indicated. Nuclear and cytoplasmic fractions were isolated and probed for the level of β-catenin. Tubulin and Lamin B1 were the controls for the cytoplasmic and nuclear proteins, respectively.

A potential caveat associated with our observations is that the established human cancer cell lines we used may have acquired mutations *in vitro* that predispose them to apoptosis in an exaggerated manner not seen with the tumors in patients. We therefore decided to test whether primary lymphoma cells isolated fresh from patients are sensitive to SR-3029. We found that SR-3029 potently killed primary lymphoma cells isolated from 3 MCL patients and 1 adult T-cell lymphoma/leukemia (ATLL) patient (**Figure 3D**), suggesting that primary lymphoma cells and cell lines alike are highly susceptible to killing by the CK1δ inhibitor SR-3029. Other blood cancers, such as multiple myeloma (MM), were also very sensitive to SR-3029 (**Figure 3E**). Our hypothesis predicts that cells with lower CK1δ protein levels will be less responsive to SR-3029. To test this idea, we assayed cell viability in peripheral blood mononuclear cells (PBMCs) which exhibited very low CK1δ protein levels compared to lymphoma cell lines (**Figure S3B**). We found that PBMCs were more resistant to SR-3029 than lymphoma cell lines (**Figure S3C Table S1)**. Our results indicate that SR-3029 can kill blood cancers in a histology-agnostic manner. Furthermore, the differential protein levels of CK1δ and correlated survival responses in the lymphoma cells versus the PBMCs suggest a potential therapeutic window for SR-3029.

SR-3029 was previously reported to inhibit tumor growth in breast cancer models via targeting Wnt/β-catenin signaling mediated by CK1δ [6]. Because dysregulated Wnt/β-catenin is highly prevalent in colon cancer, we were interested in studying the activity of SR-3029 in this context. First we confirmed that the breast cancer cell line MDA-MB-231 was sensitive to SR-3029 (**Figure S3D)**. We next tested SR-3029 in colon cancer using patient-derived tumor organoids, which have the advantages of avoiding *in vitro* acquired genetic alterations in established cell lines and recapitulating parental tumor heterogeneity in three-dimensional culture[13]. We observed that SR-3029 potently inhibited cancer cell viability of various colon cancer organoids with IC_50_ values ranging from 60 to 371 nM (**Figure S3E**). We next wished to determine if SR-3029 kills solid tumors regardless of histology. We found that the human pancreatic ductal adenocarcinoma (PDAC) cell lines (MiaPaca-2 & ASPC-1) were significantly more resistant to SR-3029 than the solid and liquid tumor models described above (**Figure S3F**). These results demonstrate that SR-3029 is toxic to select solid tumors that are possibly dependent on CK1δ-mediated signaling pathways.

Lymphomas are a highly heterogeneous group of blood cancers. Even within DLBCL multiple subtypes are defined by distinct mutations, gene expression profiles, treatment responses, and prognosis [14, 15]. We reasoned that the broad cytotoxicity of SR-3029 to lymphomas indicates that its mechanistic target is unlikely to be restricted to a molecular subtype such as those defined by MYD88 and BCL6 [14, 15]. Rather, fast growing lymphomas may be commonly dependent on activated translation associated with hyperphosphorylated 4E-BP1. We found that both 4E-BP1 and RPS6 were highly phosphorylated in all four MCL cell lines that we studied, and SR-3029 treatment for 24 hours potently inhibited phosphorylation of both 4E-BP1 and RPS6 (**Figure 2C, 3F-H**). To determine whether the above mechanism of SR-3029 in MCL is broadly applicable in lymphomas, we next studied DLBCL, the most common aggressive lymphoma. SR-3029 potently inhibited phosphorylation of 4E-BP1 and RPS in the DLBCL cell line OCI-LY10 and to a lesser degree in the DLBCL cell line OCI-LY7 (**Figure 3I, 3J**). These results indicate that 4E-BP1 is a conserved target of SR-3029 across lymphomas.

Because SR-3029 has been reported to act through inhibiting nuclear localization of β-catenin in breast cancer [6], an intriguing question is whether SR-3029 also modulates 4E-BP1 in breast cancer. In the human breast cancer cell line MDA-MB-231 phosphorylation of 4E-BP1 was only mildly reduced by SR-3029 at the concentration range of 100-300 nM with 20h exposure, and phosphorylation of RPS6 was not at all affected by SR-3029 (**Figure S3G**). We confirmed that SR-3029 inhibited nuclear localization of β-catenin in MDA-MB-231 (**Figure S3H**), as previously described [6]. In contrast, SR-3029 did not significantly reduce nuclear localization of β-catenin in the MCL cell line Z138 and DLBCL cell line OCI-LY10 at concentrations that potently reduced phosphorylation of 4E-BP1 and RPS6 (**Figure 3K, 3L**). These results suggest that the critical targets of SR-3029 in liquid tumors are regulators of translation initiation such as 4E-BP1, whereas in solid tumors SR-3029 targets alternative pathways such as Wnt/β-catenin.

### SR-3029 and mTOR inhibitors differentially inhibit tumor growth and 4E-BP1 phosphorylation

Because SR-3029 and mTOR inhibitors appear to share the same mechanistic target, namely 4E-BP1 phosphorylation, clinical experiences with mTOR inhibitors may be useful to guide the future development of CK1δ inhibitors. We therefore compared the cytotoxic and molecular effects of SR-3029 to mTOR inhibitors including Torin-1 and Everolimus. Remarkably, we found that SR-3029 was substantially more cytotoxic than Torin-1 in several lymphoma models. In the DLBCL cell line OCI-LY10, SR-3029 potently reduced cell viability in a concentration dependent manner, reducing viability to near 0% at 200 nM; in contrast, Torin-1 reduced the viability of OCI-LY10 to near 50% at 50 nM and the inhibition then plateaued at the concentration range of 50-500 nM (**Figure 4A**). After treatment with SR-3029 for 24h in OCI-LY10 cells, we observed marked induction of PARP cleavage. In contrast, Torin-1 and Everolimus induced little or no PARP cleavage (**Figure 4B**). Similarly, SR-3029 was significantly superior to Torin-1 in the MCL lymphoma cell Z-138 (**Figure 4C-D**) and the primary lymphoma cells isolated from a patient with the leukemia subtype of ATLL (**Figure 4E**). We noted that in some lymphoma cell lines such as the DLBCL cell line OCI-LY7 and MCL cell line Jeko-1, the cytotoxic effect of Torin-1 did not level off and was mildly more potent than SR-3029 (**Figure S4A-B**). The DLBCL cell line VAL, which is characterized by a complete absence of 4E-BP1 protein, was resistant to both Torin-1 and SR-3029 (**Figure S4C**).

**Figure 4.**
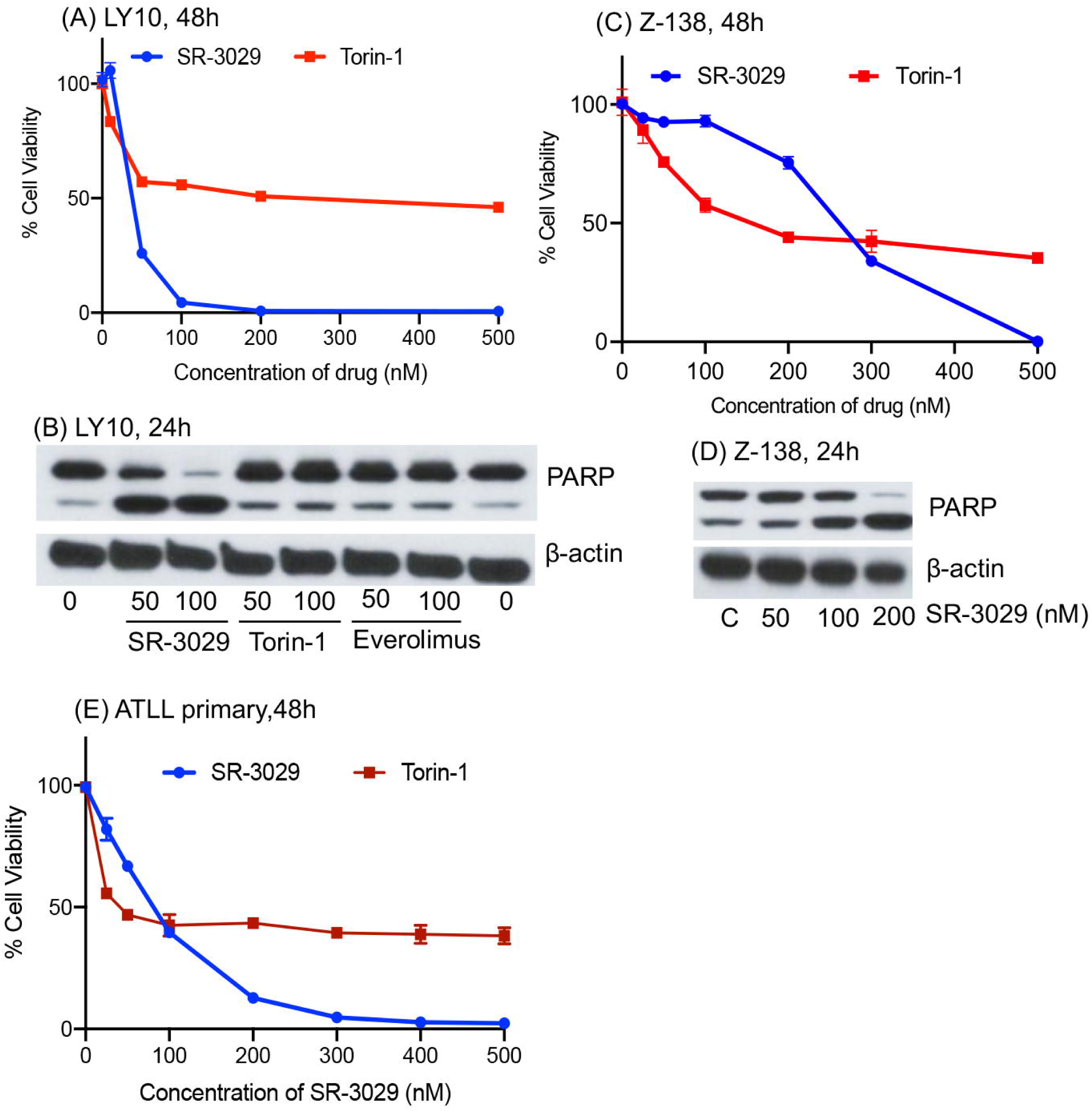
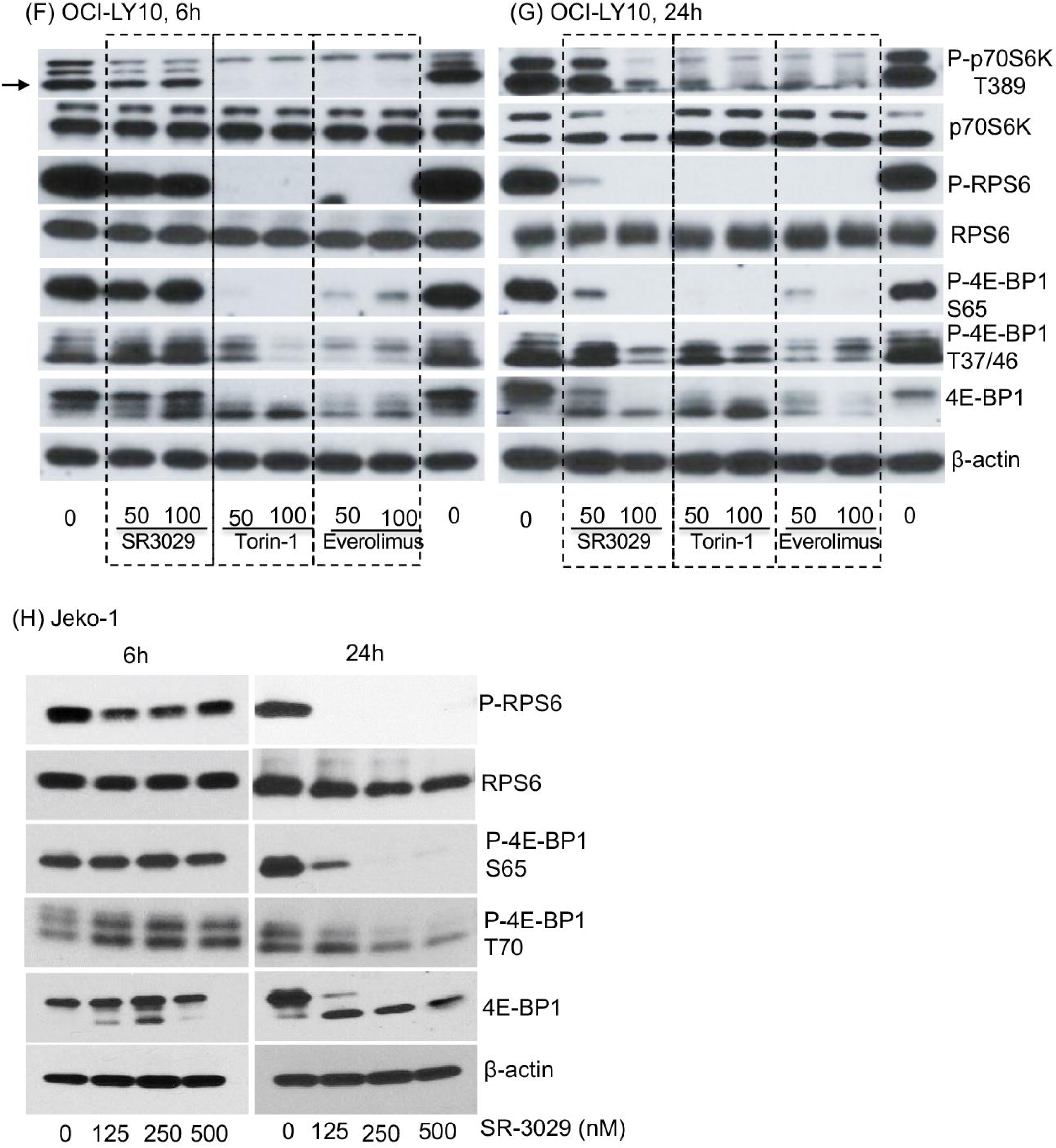
SR-3029 and mTOR inhibitors differentially inhibit tumor growth and 4E-BP1 and p70S6K phosphorylation. **(A-B)** OCI-LY10 cells treated with SR-3029, Torin-1, and the negative control DMSO. Survival cells after 48h treatment were quantitated by the Cell Titer Glo assay (A). PARP cleavage was determined by immunoblotting in cells treated for 24h (B). **(C-D)** Z-138 cell line was treated with SR-3029, Torin-1, and the negative control DMSO. The survival curves were generated from the data of 48h treatment (C). Immunoblot was performed using the samples treated for 24h (D). **(E)** Primary ATLL cells were isolated from a patient and treated with SR-3029, Torin-1, and the negative control DMSO for 24 and 48h, and the viable cells were quantitated by the Cell Titer Glo assay. The survival curves were generated from the data of 48h treatment. **(F-G)** OCI-LY10 cells were treated as indicated for 6 and 24h and analyzed by Immunoblotting. Arrows indicate the specific band of phospho-p70S6K. **(H)** Jeko-1 cells were treated as indicated for 6 and 24h and analyzed by Immunoblotting.

The superior anti-lymphoma activity of SR-3029 relative to Torin-1 led us to investigate whether they differentially affect the substrates of mTORC1 (4E-BP1 and p70S6K) and mTORC2 (AKT). Surprisingly, we found SR-3029, the more cytotoxic agent, was substantially inferior to Torin-1 and Everolimus in downregulating phosphorylation of 4E-BP1 and p70S6K/RPS6 at the short treatment time of 6 hours in OCI-LY10 cells (**Figure 4F**). Interestingly, as the treatment continued to 24h, SR-3029 exhibited nearly complete inhibition of the above phosphoproteins in a manner comparable to Torin-1 and Everolimus in OCI-LY10 (**Figure 4G**). Both SR-3029 and Torin, at 100 nM and 24h, abolished phosphorylation of AKT-S473, although at 6h they induced minimal and mild inhibition of phospho-AKT-S473, respectively (**Figure S4D-E**). In a similar manner we observed that a relatively long treatment time of 24h (but not 6h) was necessary and sufficient for SR-3029 to potently inhibit phosphorylation of 4E-BP1 in other lymphoma models, including MCL lymphoma Jeko-1 and Z-138 (**Figure 4H, 2C, S4F**) and DLBCL lymphoma OCI-LY7 (**Figure S4G-H**). Phosphorylation of p70S6K/RPS6 was highly responsive to 24h treatment of SR-3029 in the MCL cell lines Z-138 (**Figure 2C**) and Jeko-1 (**Figure 4H**), but only mildly or minimally responsive in the DLBCL cell lines OCI-LY7 (**Figure S4G-H**) and VAL (**Figure S4I-J**).

In contrast, phosphorylation of p70S6K/RPS6 was highly responsive to Torin-1 at 6h and 24h in OCI-LY7 (**Figure S4G-H**) and VAL (**Figure S4I-J**). In the MCL cell lines Jeko-1 and Z-138 phosphorylation of RPS6 was potently inhibited by Everolimus and Torin-1 at 24h, and 4E-BP1 phosphorylation was markedly repressed only by Torin-1 but not by Everolimus (**Figure S4K-L**).

These results indicate that in lymphoma cells SR-3029 potently inhibits 4E-BP1 phosphorylation but with delayed kinetics compared to mTOR inhibitors (24h vs. 6h). The activity of SR-3029 to inhibit p70S6K/RPS6 phosphorylation ranges from minimal to potent depending on the lymphoma cell lines.

### SR-3029 inhibits polysome formation and global protein synthesis

Given the effects of SR-3029 on eIF4F we predicted that SR-3029 will interfere with mRNA translation. We first examined the effects of SR-3029 on global translation by analyzing polysome profiles. In the Z-138 lymphoma cells treated with the DMSO vehicle control the 40S, 60S, 80S ribosome peaks were easily identified and the polysome levels were high (**Figure 5A**). RPS6 was absent from the RNP fractions, but was present throughout the 40S, 60S, 80S, and polysome fractions with enrichment in the 80S and heavy polysomes (**Figure 5A**). β-actin should not associate with ribosomes and as such was present only in the RNP fractions but absent from the ribosome and polysome fractions (**Figure 5A**). CK1δ and eIF4E were both present in the RNP fractions as expected but were also abundant in the ribosome fractions which is consistent with their roles as cap-associated proteins (**Figure 5A**). SR-3029 treatment resulted in a substantial increase in the 80S monosome peak with a concomitant decrease of polysomes clearly indicating that this compound inhibits global translation (**Figure 5B**). Accordingly, RPS6 became concentrated in the 80S fraction and absent from the polysome fractions (**Figure 5B**). Furthermore, SR-3029 treatment caused both CK1δ and eIF4E redistribution from the polysomes to the 40S fraction (**Figure 5B**). In comparison, Everolimus produced a weaker repression of global translation compared to SR-3029 in Z-138 cells (**Figure 5C**). Global repression of translation in response to Torin-1 was similar to SR-3029 (**Figure 5D**).

**Figure 5.**
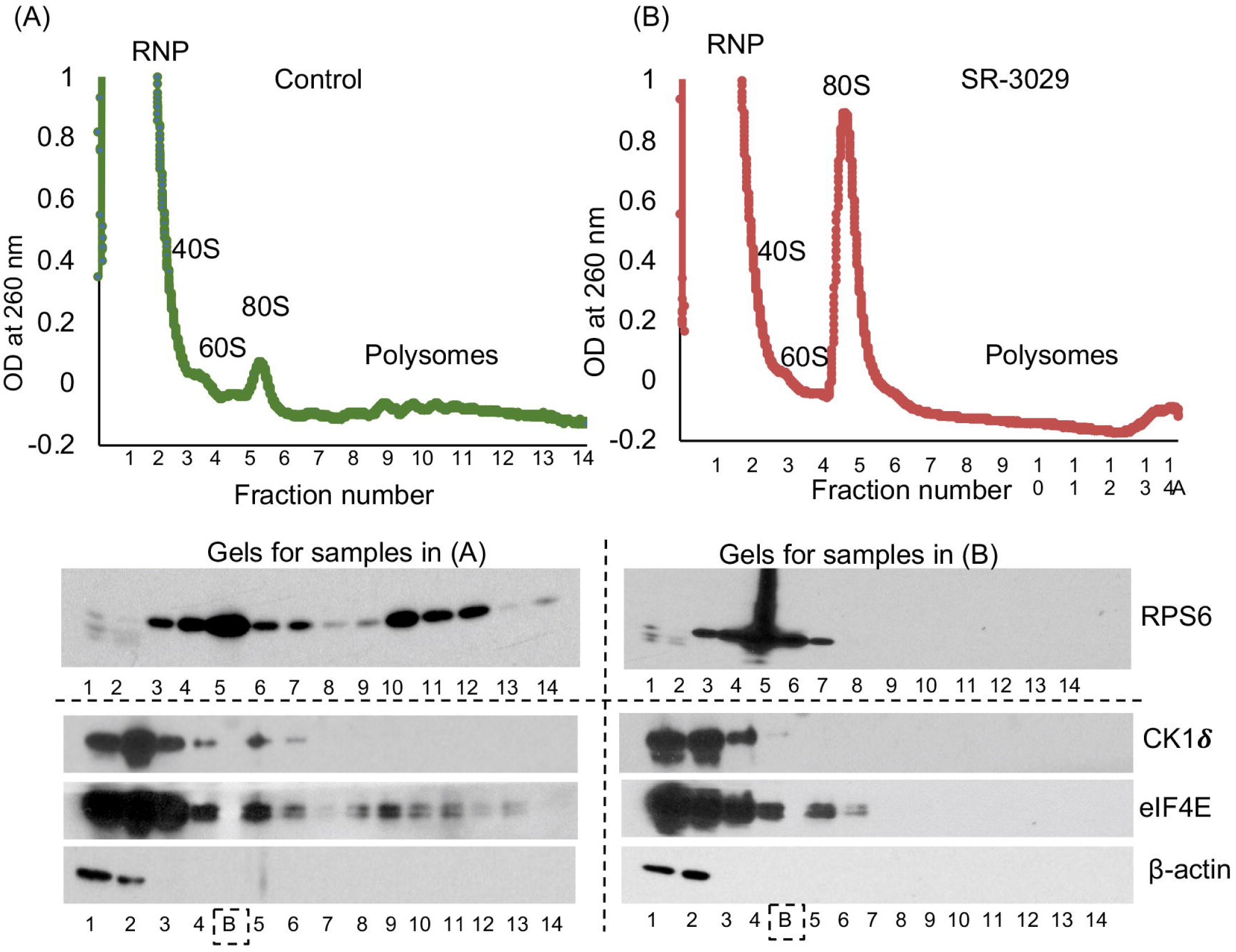
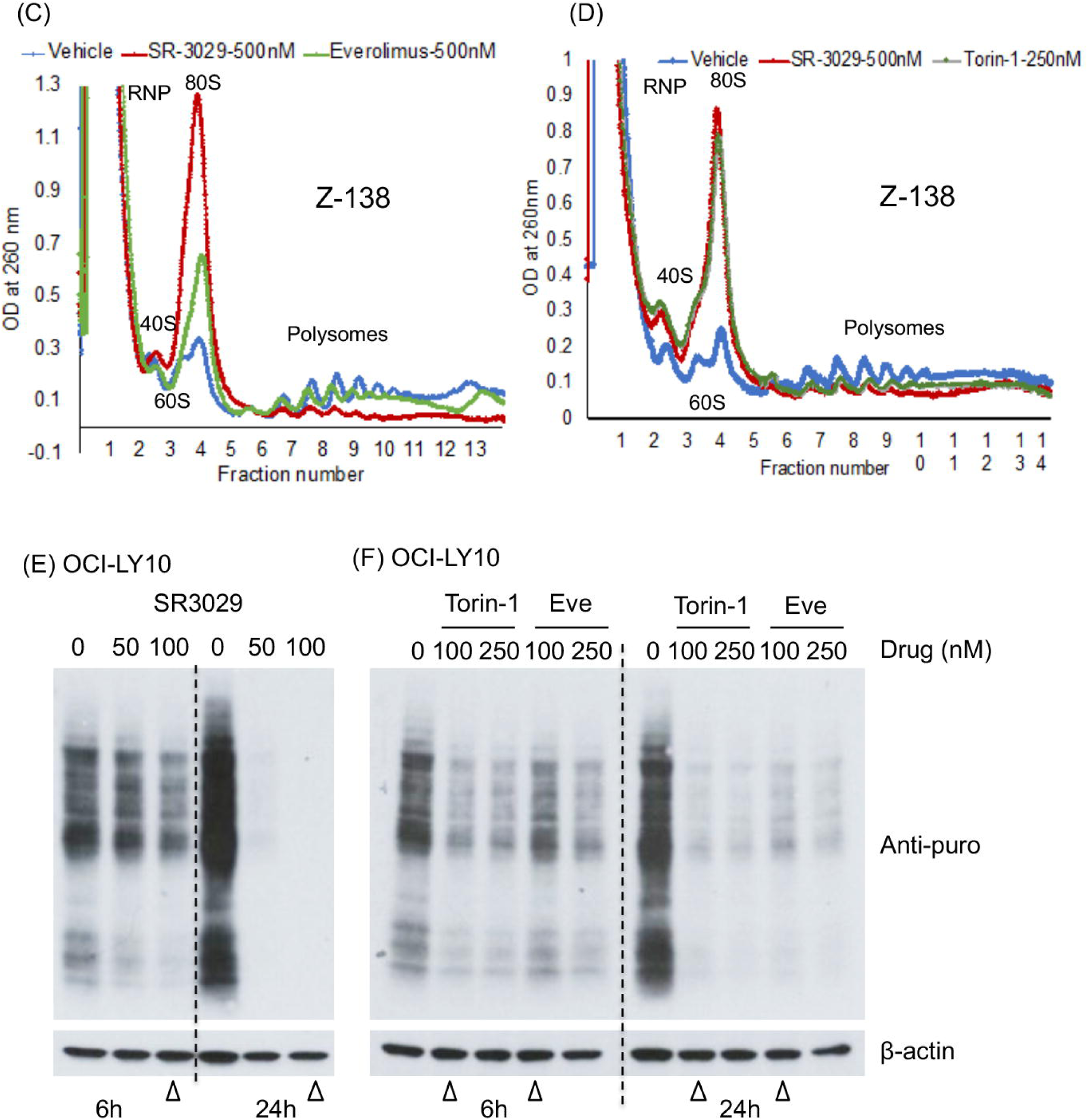
SR-3029 inhibits polysome formation and global protein synthesis. **(A-B)** Polysome profiling and immunoblotting of the fractions. The Z-138 cells were treated with DMSO vehicle control (A) or SR-3029 500 nM (B) for 6h. The cell lysates were fractionated by 10–50% sucrose density gradient centrifugation. The levels of mRNA in the sedimentations (fractions) were measured by OD260 UV recording. Polysome profiles were shown on the top graph, and data shown were the average of three experiments. In addition, the fractions of each sample were run on two gels and immunoblotting was done using the indicated antibodies. The upper gels were run from Fraction 1 to 14 without interruption, and the lower gels were run with an empty lane between Fraction 4 and 5 (indicated by “B” and an empty square). The immunoblotting was repeated twice. RNP, ribonuclear protein; 40S, 40S small ribosome subunit; 60S, 60S large ribosome subunit; 80S, 80S monosome. **(C-D)** Polysome profiling. The Z-138 cells were treated with DMSO vehicle, SR-3029 500 nM, Everolimus 500 nM, or Torin-1 at 250 nM for 6h. The cell lysates were fractionated by 10–50% sucrose density gradient centrifugation. The levels of mRNA in the sedimentations (fractions) were measured by OD260 UV recording. Data shown were the average of three experiments. **(E-F)** OCI-LY10 cells were treated with SR-3029 (SR), Torin, Everolimus (Eve) at the indicated concentrations or the DMSO control for 6h or 24h. Puromycin was then added at the final concentration of 1μg/ml for 30 min. Cell lysates were subjected to Western blotting using the anti-puromycin or anti-β-actin antibody. The empty triangles point to samples treated with 100 nM of the indicated drugs.

To further understand how SR-3029 influences global protein synthesis we utilized the SUnSET assay as described above. In the DLBCL cell line OCI-LY10, SR-3029 potently inhibited global protein synthesis in a concentration (50-100 nM) and time (6-24h) dependent manner (**Figure 5E**), which correlated with the changes in 4E-BP1 and RPS6 phosphorylation (**Figure S5A**). Furthermore, with 24h exposure SR-3029 was more potent than Torin-1 and Everolimus in blocking protein synthesis (**Figure 5E-5F**). Similarly, SR-3029 at 24h almost completely blocked translation in the MCL cell lines Jeko-1 and Maver-1 (**Figure S5B, S5C**), and SR-3029 was more potent than Torin-1 and Everolimus when assayed at 24h in the MCL cell line Z-138 (**Figure S5D, S5E**). In contrast, in the breast cancer cell line MDA-MB-231 SR-3029 at 250 nM and 24h produced no inhibition, and at 500 nM and 24hours, SR-3029 produced only a mild inhibition of protein synthesis (**Figure S5F**). This lack of response to SR-3029 in protein synthesis in MDA-MB-231 stands in stark contrast to how effectively SR-3029 inhibits the breast cancer cell viability and nuclear localization of β-catenin described above. These results demonstrate that in lymphoma cells SR-3029 given over 24h effectively shuts down global protein synthesis, and it tends to be more effective than the mTOR inhibitors. In contrast, even when a solid tumor (e.g. MDA-MB-231) is sensitive to SR-3029 the drug may not effectively inhibit global protein synthesis.

### SR-3029 causes translational downregulation of translation-related genes and cancer driver genes

Our results have so far indicated that SR-3029 inhibits translation initiation and protein synthesis. To determine whether and how SR-3029 modulates specific mRNA and protein levels, we conducted RNAseq and whole-proteome quantitative mass spectrometry in two lymphoma cell lines. Our hypothesis was that if SR-3029 specifically inhibits translation, then genes that are regulated primarily at the translation step (vs. transcription) will have reduced protein levels that cannot be accounted for by decreases of their mRNA levels. The lymphoma cells were treated with DMSO control or SR-3029 for 3 hours, and the short exposure time was chosen to minimize secondary non-specific changes of gene expression that can happen with extended treatment course. We performed pathway enrichment analysis on the SR-3029 induced signatures of change in mRNA and protein expression, using the MSigDB collections [16] gene ontology (N=9,996 gene sets) and Reactome (N=1,499 curated gene sets). The most down- and upregulated pathways were ranked by Benjamini-Hochberg FDR-corrected p-value. Strikingly, 7 of the 8 most down-regulated gene ontology pathways in the protein expression signature of SR-3029 directly involved translation initiation, ribosome, and or co-translational targeting, all with FDR p-value <10^−8^ (**Figure 6B**). Conversely, the opposite effect was seen in the mRNA signature, where there was marked upregulation in these same translation initiation pathways (**Figure 6A**). SR-3029 consistently demonstrated a selective and marked effect on eukaryotic translation initiation and early co-translational pathways, i.e. significant decrease in the protein and simultaneous increase in mRNA levels, across treatment conditions in two different lymphoma cell lines (**Figure 6A, 6B**; **Figure S6A, S6B**). The discordance in upregulated mRNA and downregulated protein levels was salient for the 40S ribosomal genes (**Figure 6C**). In contrast, there were concordant decreases in the mRNA and protein levels among the post-translational modification pathway genes (**Figure 6C**). Despite the inverse relationship between RNA and protein levels for these pathways/genes discussed above, at the global level RNAseq and proteomics data exhibited a near total decoupling of mRNA and protein expression, with the correlation of the signatures being zero in all tested conditions (**Figure 6D**). These results indicate that many translation-related genes are among the first to be reduced specifically at the translation step by a short treatment of SR-3029.

**Figure 6.**
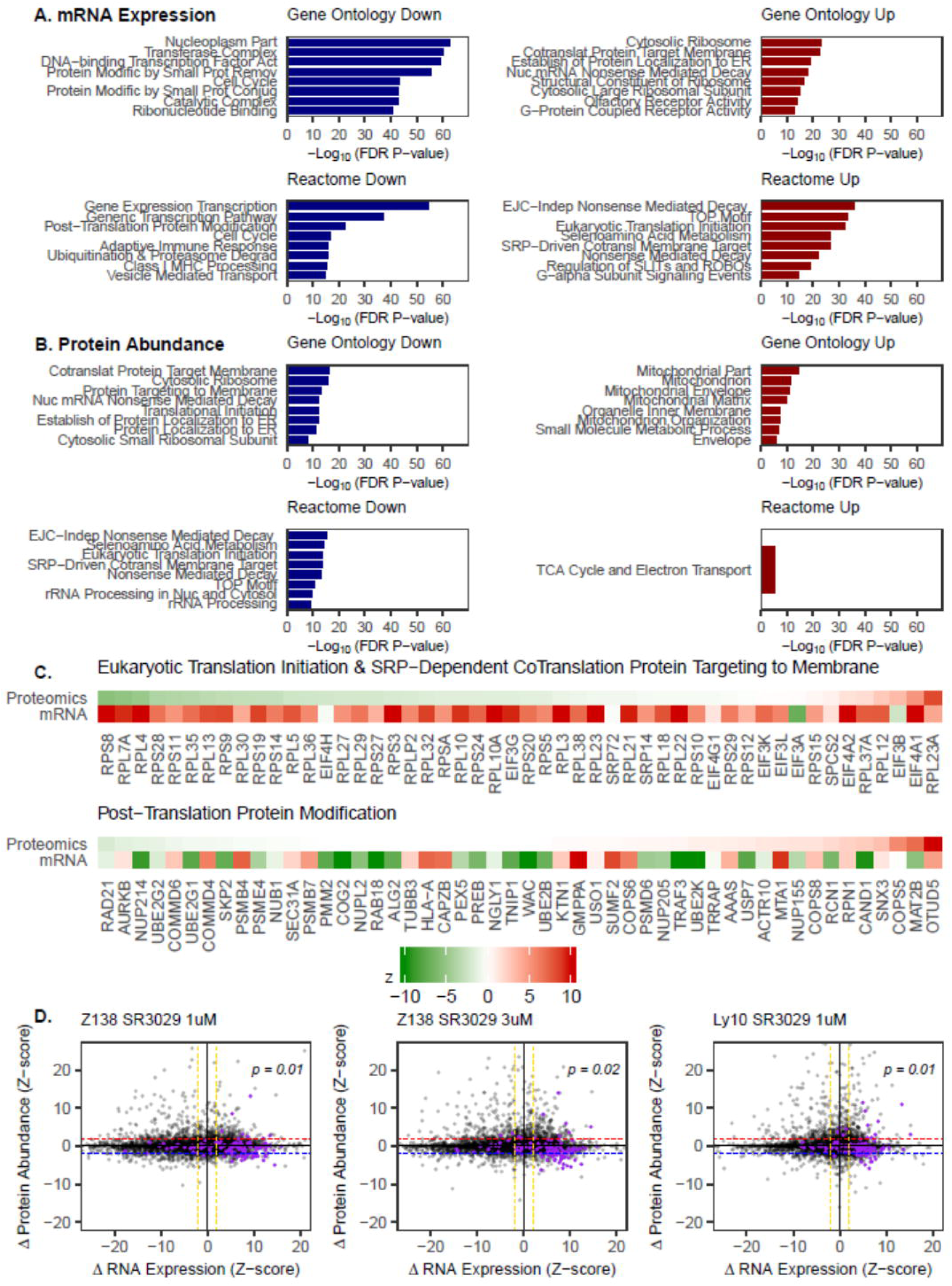
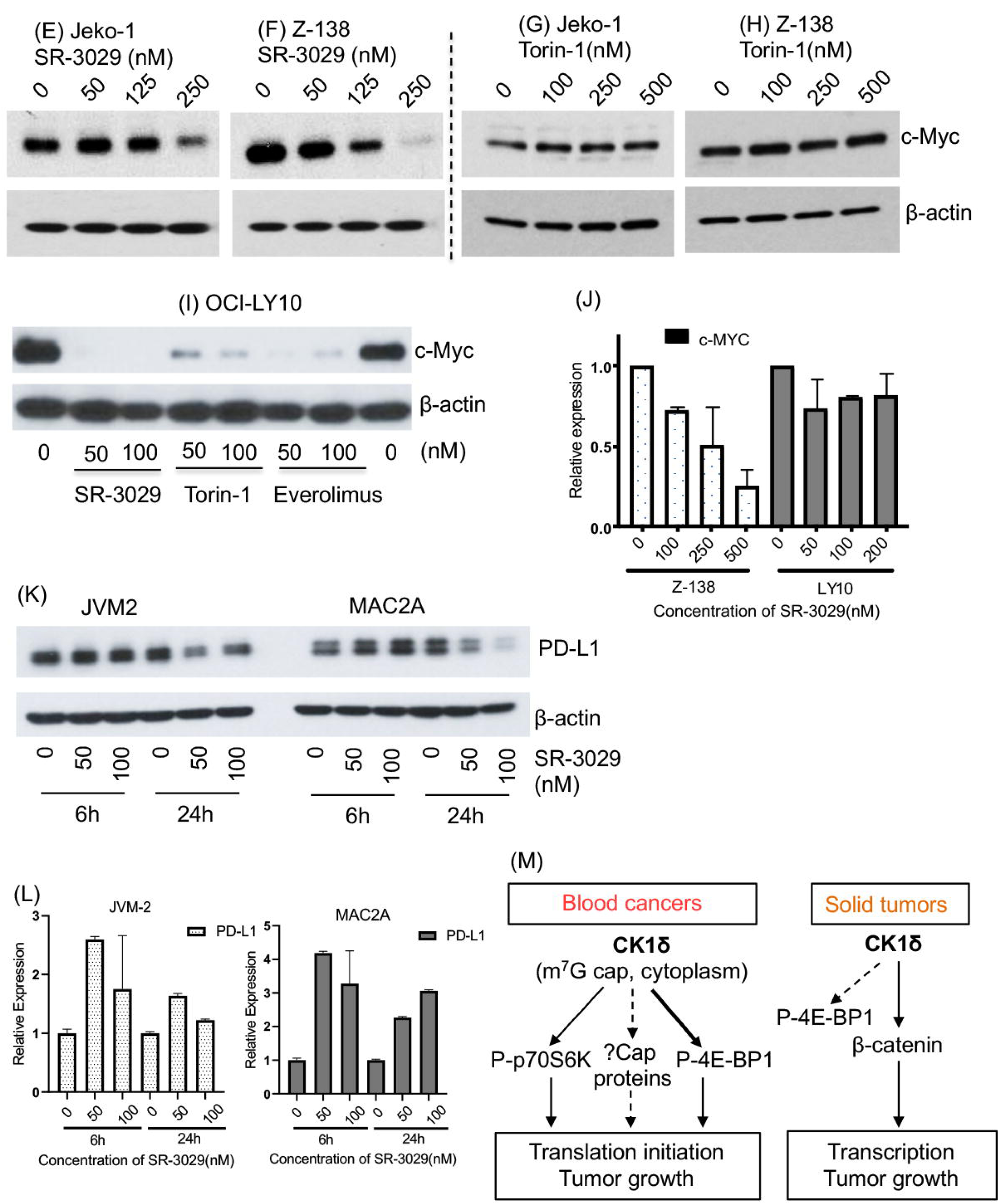
SR-3029 downregulates global gene expression and inhibits translation of C-MYC and PD-L1. **(A-B)** Z-138 lymphoma cells were treated with 3 *µ*M SR-3029 or DMSO control for 3h, and processed for RNAseq and proteomics studies. The RNA and protein abundance of each gene were compared between the treatment and control samples. Pathway analysis was performed on the SR-3029 induced differential RNA expression signature (A) and the proteomics expression signature (B) using the analytic rank-based enrichment analysis algorithm (a version of preranked GSEA). The most enriched Gene Ontology pathways and Reactome pathways are shown, ranked by FDR-corrected p-value, with down and up pathways displayed separately. **(C)** Upper panel: Heatmap of the top 50 most differentially expressed genes/proteins involved in the Reactome Eukaryotic Translation Initiation pathway [R-HSA-72613] and SRP-dependent cotranslational targeting of proteins to ER/cell membrane [R-HSA-1799339]. SR-3029 decreased the abundance of 40S ribosomal proteins but increased the mRNA expression of these genes, consistent with a failure of a stable translation initiation complex and polysome formation. Lower panel: Heatmap of the top 50 most differentially expressed genes/proteins in the Reactome post-translational modification pathway [R-HSA-597592]. At the mRNA level, many genes involved in post-translational modifications are underexpressed, which may be a consequence of cessation of protein translation. **(D)** Scatter plots of the protein expression signature vs. mRNA expression signature [SR-3029 vs DMSO control]. Three conditions are shown as analyzed. There is a complete decoupling of mRNA and protein expression, with zero correlation in the signatures. The purple points highlight genes involved in translation initiation and are over-represented in the lower right quadrants with decreased protein abundance and increased mRNA expression. **(E-L)** The specified lymphoma cell lines were treated with SR-3029, Torin, Everolimus at the indicated concentrations or the DMSO control. The treatment was 24h unless specified as 6h in (J, K, L). The samples were analyzed by immunoblot unless specified as qPCR in (J, L). **(M)** Models of CK1δ in blood cancers and solid tumors. Dashed arrows indicate where the regulation is weak or hypothetical.

We chose to focus our investigation on C-MYC and PD-L1 because they have therapeutic and prognostic significance in many cancers, and their translation is repressed by translation inhibitors [17-19]. We found that the protein level of C-MYC in the MCL cell lines Jeko-1 and Z-138 was moderately or near completely repressed by SR-3029 at 24h (**Figure 6E-F**). In contrast Torin-1, even at 500 nM, failed to reduce C-MYC protein level (**Figure 6G-H**). In the DLBCL cell line OCI-LY10, SR-3029 completely turned off the C-MYC protein while Torin-1 and Everolimus still allowed some C-MYC protein to be synthesized (**Figure 6I**). We next wanted to test whether the SR-3079-dependent repression of C-MYC protein is associated with downregulated C-MYC transcription. In OCI-LY10 the mRNA level of C-MYC remained constant with the SR-3029 treatment (**Figure 6J**). Thus, the repression of C-MYC protein by SR-3029 treatment in OCI-LY10 is predominantly post-transcriptional. Based on the above results strongly implicating CK1δ and SR-3079 in translation we favor the hypothesis that SR-3079 causes translational repression of C-MYC in the DLBCL cell line OCI-LY10. In the MCL cell line Z-138 the mRNA level of C-MYC was moderately repressed by SR-3029 (**Figure 6J**), suggesting that in Z-138 SR-3029 may decrease the protein level of C-MYC through transcriptional and translational mechanisms. We next assayed PD-L1 in the MCL cell line JVM2 and PTCL cell line MAC2A, because they were the only two among twelve unselected lymphoma cell lines that had a high level of PD-L1 protein (**Figure S6C**). We found that SR-3029 treatment decreased the protein level of PD-L1 mildly in JVM2 and substantially in MAC2A (**Figure 6K**), but did not repress the mRNA level of PD-L1 in either of these cell lines (**Figure 6L**). These results suggest that SR-3079 can inhibit translation of critical cancer drivers such as C-MYC and PD-L1 in select lymphoma models.

## DISCUSSION

Here we present evidence that CK1δ is a critical regulator of eIF4F-dependent translation initiation. We show that the involvement of CK1δ in translation initiation is essential in blood cancers but is limited in solid tumors (e.g. breast cancer, pancreatic cancer) and non-neoplastic cells (e.g. PBMC, HEK293). The mechanism of CK1δ in regulating translation in blood cancers may be twofold (**Figure 6M**): first, CK1δ clearly regulates phosphorylation of 4E-BP1, which is a direct substrate of mTOR in its regulation of translation initiation. CK1δ also regulates phosphorylation of p70S6K, however, this effect is less universal but is variable in different lymphoma cell lines. The role of CK1δ in these phosphorylation events is more likely indirect, thus distinct from the direct effect of mTOR. Second, CK1δ regulates eIF4F, and a substantial proportion of CK1δ is localized to the mRNA m^7^G cap, which is similar to eIF4E. CK1δ does not co-immunoprecipitate with eIF4E, suggesting the localization of CK1δ at the cap is unlikely mediated by its binding to eIF4E but more likely through its interaction with other cap-associated proteins. Interestingly, a quantitative proteomic study of mRNA cap in HEK293 does not identify CK1δ among more than 160 cap-associated proteins [12]. Our results indicate that CK1δ may be a critical kinase at the mRNA cap, where CK1δ may phosphorylate translation regulators to facilitate translation initiation. Finding the protein substrates of CK1δ at the m^7^G cap will provide new insights into translation regulation and potential therapeutic targets in blood cancers.

A number of compounds that target eIF4F have demonstrated potent anti-tumor activities. Unfortunately, their progress in the clinic has been disappointing. Compared to eIF4A, which has been extensively studied as a therapeutic target, CK1δ as a kinase and drug target offers clear practical advantages due to the prior successes of kinase inhibitors approved for cancer treatment. Targeting CK1δ using SR-3029 results in potent anti-tumor activity in lymphomas and myeloma. These lymphomas are heterogeneous in terms of histology, pathogenetic factors, treatments, and outcomes. Remarkably, SR-3029 kills nearly all lymphoma cell lines that we tested. Our data support the hypothesis that many lymphomas, regardless of histology, share a dependence on CK1δ-driven translation initiation and are highly susceptible to killing by SR-3029 with rare exceptions. In contrast, non-neoplastic cells such as PBMCs and HEK293 are resistant to SR-3029. We hypothesize that the dependence on CK1δ in lymphomas could be reflective of the high rates of global translation in these fast-growing malignancies.

SR-3029 is well tolerated in mouse models [6]. This suggests that a reasonable therapeutic window may exist for SR-3029 or its derivatives for future use in human patients. We propose that the levels of CK1δ, eIF4E, total and phospho-4E-BP1, and total and phospho-p70S6K in the pre-treatment tumor tissues should be explored as potential biomarkers for patient selection in future clinical trials of SR-3029-based translation inhibitors. Alternatively, it may be possible to use patient derived organoids or patient derived xenograft mouse models of lymphoma to predict whether a patient may be a good candidate for treatment with CK1δ inhibitors.

CK1δ and mTORC1 share overlapping downstream targets, including 4E-BP1 and p70S6K/RPS6. mTOR inhibitors have been approved for the treatment of renal cell cancer but produce limited clinical benefits in other cancers, while the toxicities are extensive [20-22]. An obvious question or concern is that whether CK1δ will be limited by the same low efficacy and high toxicity observed with mTOR inhibitors. Despite their apparent similarity, CK1δ inhibitors and mTOR inhibitors will likely have substantial difference in future human studies for several reasons. The mTOR inhibitors Everolimus and Torin-1 represses 4E-BP1 and p70S6K/RPS6 phosphorylation rapidly within 6h of treatment. In contrast, inhibition by SR-3029 only becomes evident upon longer treatment exposure (e.g. 24h) in lymphoma cells. Although SR-3029 is slower than Everolimus and Torin-1 to inhibit phosphorylation of 4E-BP1 and p70S6K/RPS6, SR-3029 appears to block protein synthesis more completely than Torin 1 and Everolimus at the same concentrations at 24h and this effect is specific to lymphoma cells. Remarkably, SR-3029 reduces lymphoma cell viability to zero. In contrast, multiple lymphoma models retain a high level of viability (40-50%) when treated with Torin-1, even at high concentrations. In some lymphoma cell lines, SR-3029 can almost completely repress the protein level of the C-MYC oncogene which is not observed with Torin-1 treatment. These results suggest that SR-3029 may exploit cancer vulnerabilities that mTOR inhibitors fail to target.

As a potential therapeutic molecule SR-3029 presents many desirable properties: (a) SR-3029 is well tolerated in mice; (b) it has demonstrated potent anti-tumor activity *in vitro* in blood cancers and solid tumors via well-defined mechanisms; (c) the two mechanistic targets of SR-3029, namely translation and Wnt signaling, are highly relevant and commonly dysregulated in many malignancies; (d) SR-3029 as a translation inhibitor may not be subject to the cross-resistance mechanisms in cancers treated with currently available drugs that do not target translation; (e) SR-3029 remains highly active in lymphoma cell lines where the mTORC1/mTORC2 inhibitor Torin-1 fails to kill. Our results support clinical development of CK1δ inhibitors for the treatment of liquid and solid tumors.

## METHODS

### Cell line and reagents

JEKO-1, Z-138, JVM-2, MAVER-1(mantle cell lymphoma), H929 and MM1S (multiple myeloma cell line), SUDHL-6 and SUDHL-10 (DLBCL), FEPD, MAC2A and MTA cell lines were grown in RPMI medium with 10% fetal calf serum. MAC2A and MTA (PTCL) were grown in RPMI medium with 10% fetal calf serum and 1% sodium pyruvate. OCI-LY7 and OCI-Ly10, VAL (DLBCL) SMZ1 (PTCL) cell lines were grown in Iscove modified Dulbecco medium with 10% fetal calf serum. Fresh medium was added every 2 to 3 days, and the cells were kept at a cell concentration of 0.1 to 1× 10^6^ /mL. MDA-MB-231 and HEK293 cell lines were grown in Dulbecco’s modified eagle’s medium with 10% fetal calf serum. Cells were incubated at 37 °C in a 5% CO2 atmosphere, 95% humidity in CO2 incubator.

### Cytotoxicity assays

Cytotoxicity was evaluated using the Cell Titer-Glo Reagent (Promega) according to the manufacturer’s manual, as reported previously [2]. Experiments were carried out in 48-well plates, with each treatment in triplicate. Samples were taken at typically 24, 48, and 72 hours after treatment. Cytotoxicity was expressed by the decreasing percentage of live cells in each treatment relative to the untreated control from the same experiment, as a function of time. IC50 and IC90 for each cell line was calculated using the Graphpad prism software.

### Gene knock out with CRISPR/CAS9 system

CRISPR-Cas9-mediated knockout was delivered by pSpCas9 (BB)-2A-GFP (PX458) (Plasmid #48138, Addgene, and Cambridge, MA, USA). Sequences of sgRNAs are listed in the following supplementary table S2. Cells were sorted by GFP selection and selected for single clones by serial dilution, Cells were plated at 1-30 cells per 96-well plate to isolate single cell derived clones. After 7-10 days for expansion, clones were screened for CRISPR-mediated deletion. Genomic DNA (gDNA) was extracted by cell lysis followed by isopropanol precipitation and validated by sequencing.

### Purification of primary lymphoma cells and normal lymphocytes

Isolation of primary lymphoma cells and normal lymphocytes as described previously [2].

### Transfection

Stable transfections of OCI-LY-7 and Z-138 cells were performed by electroporation using the Neon® Transfection System (Invitrogen). Electroporation settings were selected after 24-well optimization was performed according to the manufacturer’s protocol. 5×10^6^ Ly7 Cells and 5 ug of px459 were electroporated per 100ul reaction and incubated for 48hrs at 37°C and 5% CO2. For Z138, 5×10^6^ Cells and 10 ug px459 were electroporated per 100ul reaction and in incubated for 48hrs at 37°C and 5% CO2 in 20%n FCS containing media. The cells were pooled, counted, and maintained for future experiments. For, HEK293 cell line were seeded and treated with Lipofectamine 3000 (Thermo Scientific, Waltham, MA, USA) and 1*µ*g/ml DNA in OptiMEM reduced-serum medium (Invitrogen, Life Technologies, Carlsbad, CA, USA). Cells were incubated for 16-24 hours before GFP selection.

### Protein Extraction and Western Blotting Analysis

Cells were collected by centrifugation at 1250 rpm and then washed with ice-cold PBS twice to completely remove medium. RIPA lysis buffer (Thermo Fisher Scientific) supplemented with Halt Protease Inhibitor Cocktail (Santa Cruz Biotechnology) was added to cell pellets to extract protein. Protein concentrations in lysates were measured by Pierce BCA Protein Assay Kit (Thermo Fisher Scientific) and followed by the addition of SDS loading buffer (4X) and heated at 95°C for 5 min. Equal amount of protein samples was subjected to SDS-PAGE and transferred to a PVDF membrane (Bio-Rad). The membrane was blocked in 5% BSA/Milk at room temperature for 1 hour and incubated with appropriate antibodies at 4°C overnight. Antibodies were diluted in TBST (TBS with 0.1% Tween) with 5% BSA/ Milk. On the next day, the membranes were washed with TBST and incubated with appropriate secondary antibodies at room temperature for 1 hour. Membranes were developed using the chemiluminescence detection system from Thermo Scientific.p-4E-BP1-S65, p-4E-BP1-T70, p-4E-BP1-T37/46, 4E-BP1, β-catenin, p-p70SK, P-RPS6, p70SK, RPS6, eIF4E, eIF4G, PARP, RAPTOR, Lamin B1, β-actin were purchased from Cell Signaling Technology. CK1δ, c-Myc, Cyclin D1 were purchased from Santa cruz Biotechnology. CK1ε was purchased from Abcam. Tubulin was purchased from Sigma. Subcellular fractionation was performed using NE-PER nuclear and cytoplasmic extraction reagents from Thermo Scientific as per manufacturer’s protocol and fractions were subjected to western blotting. Lamin B1 and Tubulin were used as loading control for nuclear and cytoplasmic fractions respectively.

### RNA extraction and quantitative PCR

Cell pellets were collected and then subjected to total RNA extraction using RNeasy Plus Mini Kit (QIAGEN, Cat#74136) according to the manufacturer’s instructions and cDNA was prepared from the extracted RNA using Omniscript reverese transcriptase (QIAGEN) according to the manufacturer’s instructions. The obtained cDNA samples were diluted and used for RT-qPCR by using TaqMan Gene Expression Master Mix (Thermo Fisher scientific). We used following gene specific primers (c-Myc: Hs00153408_m1, PD-L1: Hs00204257_m1) for PCR amplification and detection on the HT Fast Real-Time PCR System (Applied Biosystems from Life Technologies). GAPDH was used as an endogenous normalization control. In all experiments, the average of three independent reactions is shown with error bars indicating standard deviation RT-qPCR data were normalized to GAPDH and was estimated using the delta-delta Ct (ΔΔCt) method.

### Co-immunoprecipitation

Cells were harvested and co-immunoprecipitation analysis was carried out by using Pierce Co-immunoprecipitation kit (Thermo Scientific) according to the manufacturer’s instructions.

### CAP binding assay

Cells were harvested and lysed in CAP lysis buffer (1% Triton-X, 100mM KCL, 0.1 M EDTA, 10% glycerol, 2mM MgCl2, 20mM HEPES, protease inhibitor cocktail mixture) and were incubated on ice for 5 min at 4 °C and centrifuged at 15000 rpm for 5 minutes. The cleared lysates were incubated with pre-washed GTP-sepharose or m7-GTP-sepharose beads (AXXORA) with an additional CAP lysis buffer and incubated for 3hrs at 4 °C with occasional shaking. The beads were washed three times and proteins associated with CAP was eluted with 4X SDS loading buffer and subjected to Western blotting.

### SUnSET assay

SUnSet assay was performed as per Manufacturer’s protocol (Kerafast). Briefly, cells were treated and incubated for 30 minutes with 1ug/ml puromycin. After incubation, cells were washed with ice-cold PBS and lysed using RIPA lysis buffer. Lysis were subjected to western blotting and probed with anti-puromycin antibody (Kerafast). β-actin was used as loading control.

### Polysome profiling

Polysome profile was carried out as indicated before [23] with slight modification. Briefly, Cells were treated with indicated drugs for 6 hours and then harvested on ice in PBS containing 100ug/ml cyclohexamide. Cells were pelleted and lysed in lysis buffer (20 mM Tris-HCl (pH 7.4), 250 mM NaCl, 15 mM MgCl_2_, 0.5% Triton X-100, 1 mM DTT, 100ug/ml cyclohexamide, and Protease inhibitor cocktail, TURBO Dnase and RNAsin). Lysates were passed with 23G needle to homogenize. Lysates were cleared, separated on a 10%–50% sucrose gradient by ultracentrifugation, and fractionated using a Piston Gradient Fractionator, Biocomp. For western blotting, cells were lysed in low-salt lysis buffer (20 mM Tris-HCl (pH 7.4), 250 mM NaCl, 10 mM MgCl2, 0.5% Triton X-100, 1 mM DTT, 100ug/ml cyclohexamide, and Protease inhibitor cocktail, TURBO Dnase and RNAsin) and proteins were precipitated from the collcted fractions with 95% ethanol and Glycoblue. The precipitated proteins were washed with 70% ethanol and heated at 95°C for 5 min. Equal amount of protein samples was subjected to SDS-PAGE and followed by Western blotting.

### TMT spectrometry analysis

Cell lysis was performed in 8 M urea-based extraction buffer and extracted protein amount was quantified using BCA assay. Lysates were reduced, alkylated, and digested with trypsin. To enable multiplexing, peptide samples were labeled with TMT-11 reagents and after labeling, all the samples were pooled together in one mix. In order to reach sufficient proteome coverage, sample complexity was reduced by separating the pooled TMT mix into 3 fractions by basic reversed-phase chromatography[24]. All these fractions were analyzed on a Thermo Q-Exactive HF mass spectrometer. Data were analyzed using the MaxQuant proteomics software package and further processed for relative quantification as described in Cheng et al. [25]. A protein expression signature was then computed between drug treated samples and batch-specific controls using the eBayes method and lmFit function from the limma package. Finally, pathway analysis was performed using the analytic rank-based enrichment analysis, a faster implementation of pre-ranked GSEA, with ‘Gene Ontology’ and ‘Reactome gene sets provided in the Broad MSigDB collections.

### Total RNA-Seq Library Preparation

Total RNA was purified from cell lysates using the RNAeasy Plus Kit (QIAGEN). RNA concentration and integrity were verified using high sensitive RNA Qubit reagent (Invitrogen) and Agilent 2100 Bioanalyzer (Agilent Technologies) respectively. Ribosomal RNA was depleted using NEB next rRNA depletion kit (NEB). Strand specific sequencing libraries were generated from rRNA depleted total RNAs using the NEBNext® Ultra Directional RNA Library Prep Kit for Illumina. RNA Seq libraries were quantified using high sensitive double stranded qubit DNA reagent and library size was verified using Agilent Bioanalyzer. Sequencing was performed on an Illumina NextSeq 500 sequencer with a read length of 75 bases [26]. Approximately 20 to 30 million demultiplexed, pass-filtered, single-end reads for each sample were obtained. After standard read alignment of RNA-Seq data by STAR [27] to the GRCh38 reference genome build and summarization of expression quantities at the gene count level, gene expression was normalized by the voom function, as implemented in the limma package on Bioconductor [28]. A gene expression signature was then computed between drug treated samples and DMSO reference using the eBayes method and lmFit function from the limma package. Finally, pathway analysis was performed using the analytic rank-based enrichment analysis [29], a faster implementation of pre-ranked GSEA, with ‘Gene Ontology’ and ‘Reactome’ gene sets provided in the Broad MSigDB collections.

### Statistical significance

In vitro quantitative tests such as cytotoxicity experiments were conducted in triplicate, and the mean and standard error was calculated. Unpaired Student’s t test was used to compare two independent groups. One-way analysis of variance (ANOVA) was used when three or more independent groups were compared. All tests were two-sided. A P value of <0.05 was considered statistically significant.

## ACKNOWLEDGEMENT

CD has received research funding from: TG Therapeutics, Bayer, Amgen, Columbia University TRx Award, Columbia University CTO Pilot Award, and Columbia University Precision Medicine Pilot Award. CD is grateful of the philanthropic support from: JK, RXH, and JXM. This publication was supported in part through the National Cancer Institute Cancer Center Support Grant P30CA013696 and by the National Center for Advancing Translational Sciences, National Institutes of Health, through Grant Number UL1TR001873. The content is solely the responsibility of the authors and does not necessarily represent the official views of the NIH. We thank Drs. John Blenis and Lewis C. Cantley for critical reviews.

