## Supplementary Figures for "CK1 Delta Is an mRNA Cap-Associated Protein That Drives Translation Initiation and Tumor Growth"

### Slide 1
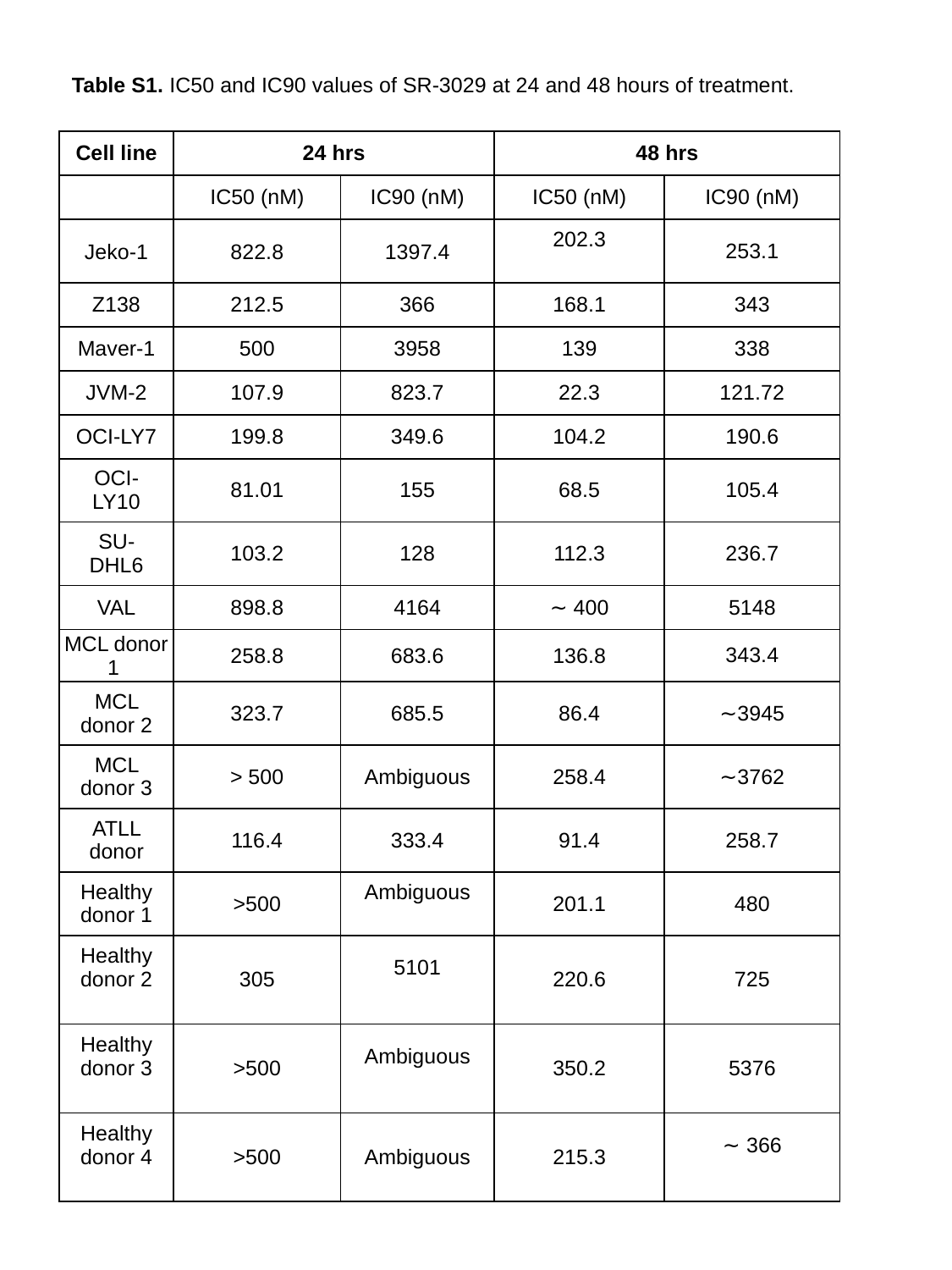

Table S1. IC50 and IC90 values of SR-3029 at 24 and 48 hours of treatment.
| Cell line | 24 hrs | | 48 hrs | |
| --- | --- | --- | --- | --- |
| | IC50 (nM) | IC90 (nM) | IC50 (nM) | IC90 (nM) |
| Jeko-1 | 822.8 | 1397.4 | 202.3 | 253.1 |
| Z138 | 212.5 | 366 | 168.1 | 343 |
| Maver-1 | 500 | 3958 | 139 | 338 |
| JVM-2 | 107.9 | 823.7 | 22.3 | 121.72 |
| OCI-LY7 | 199.8 | 349.6 | 104.2 | 190.6 |
| OCI-LY10 | 81.01 | 155 | 68.5 | 105.4 |
| SU-DHL6 | 103.2 | 128 | 112.3 | 236.7 |
| VAL | 898.8 | 4164 | ∼ 400 | 5148 |
| MCL donor 1 | 258.8 | 683.6 | 136.8 | 343.4 |
| MCL donor 2 | 323.7 | 685.5 | 86.4 | ∼3945 |
| MCL donor 3 | > 500 | Ambiguous | 258.4 | ∼3762 |
| ATLL donor | 116.4 | 333.4 | 91.4 | 258.7 |
| Healthy donor 1 | >500 | Ambiguous | 201.1 | 480 |
| Healthy donor 2 | 305 | 5101 | 220.6 | 725 |
| Healthy donor 3 | >500 | Ambiguous | 350.2 | 5376 |
| Healthy donor 4 | >500 | Ambiguous | 215.3 | ∼ 366 |

### Slide 2
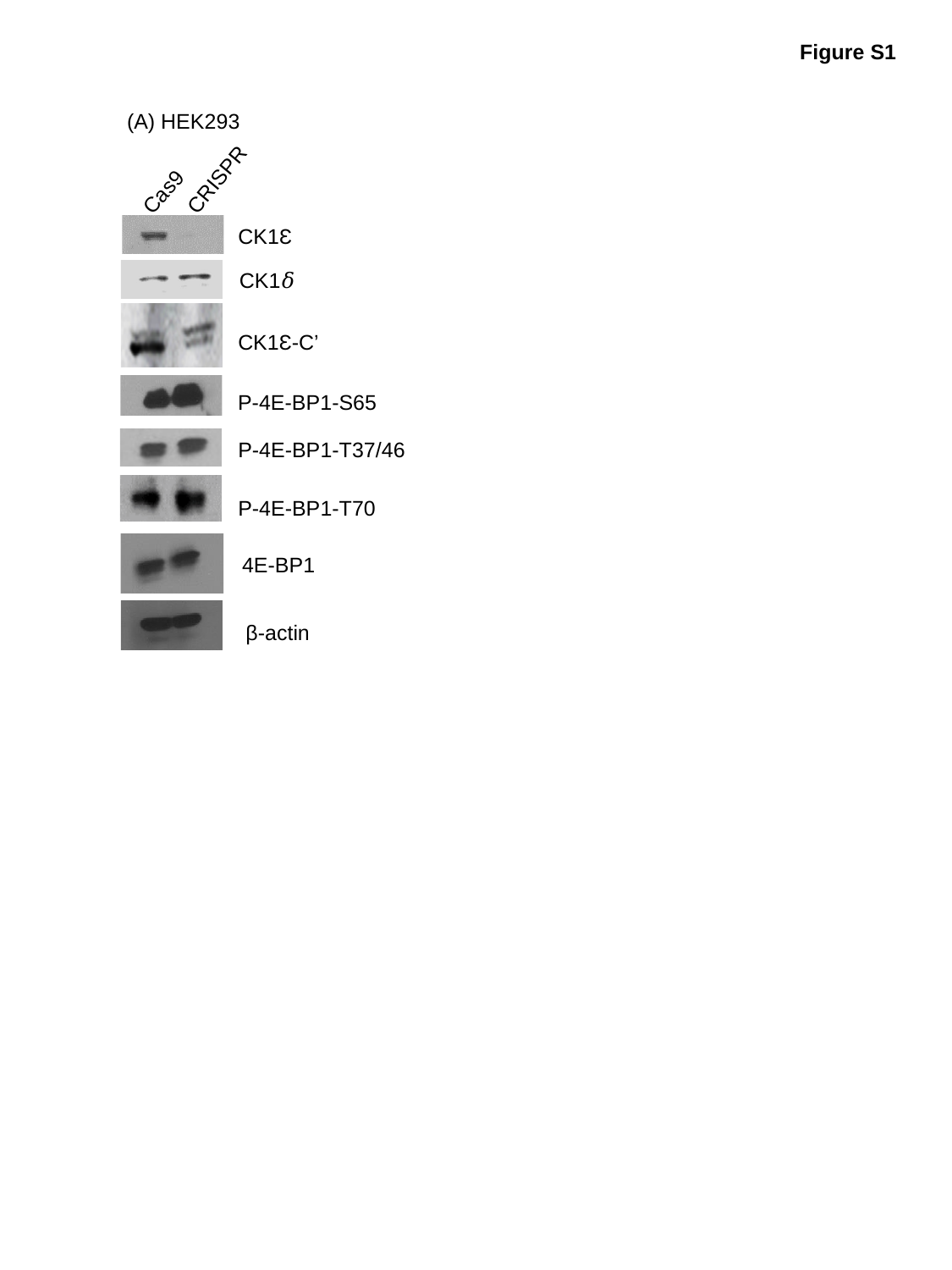

Figure S1
(A) HEK293
CRISPR
Cas9
CK1Ɛ
CK1𝛿
CK1Ɛ-C’
P-4E-BP1-S65
P-4E-BP1-T37/46
P-4E-BP1-T70
4E-BP1
β-actin

### Slide 3
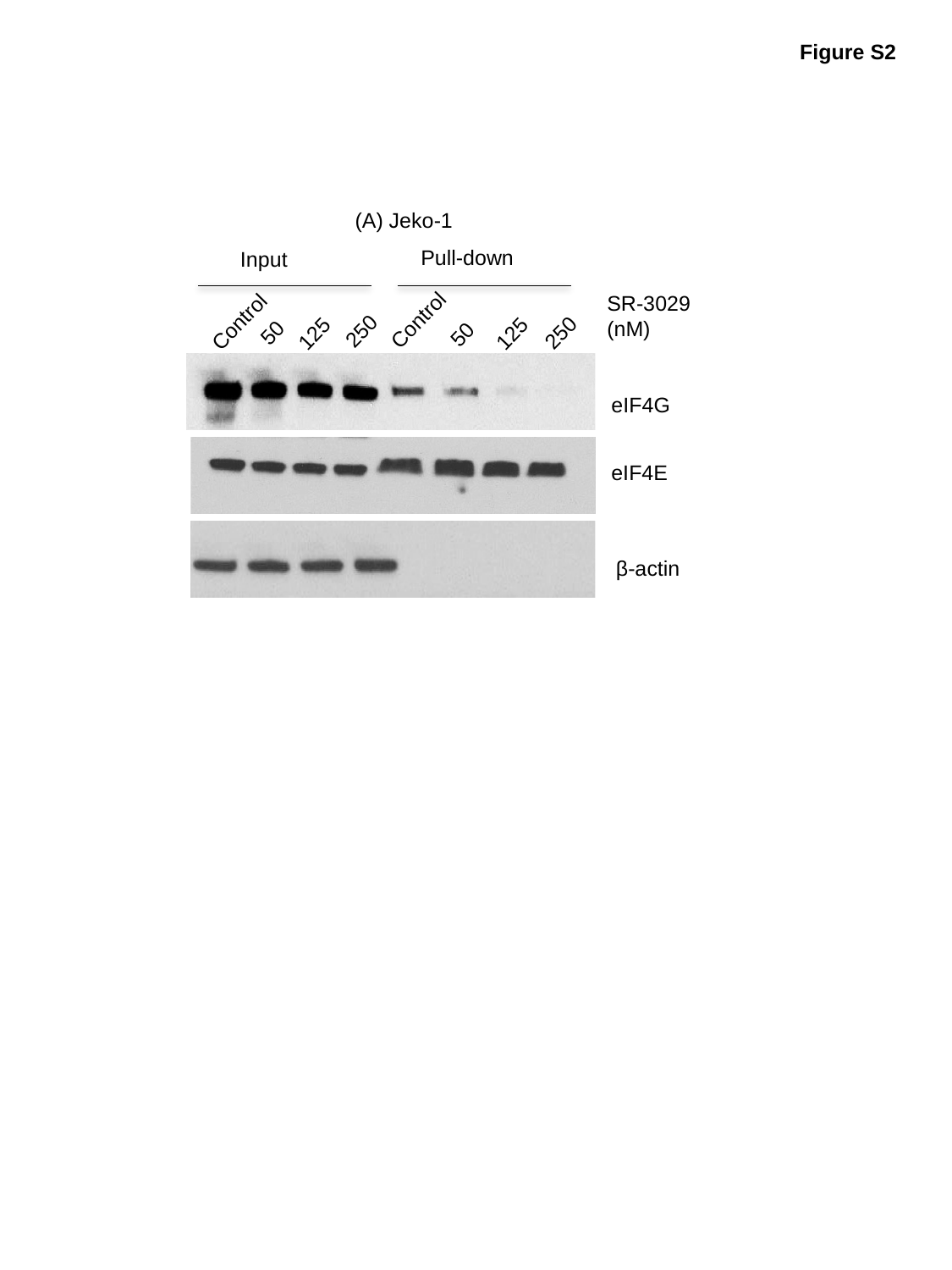

Figure S2
(A) Jeko-1
Pull-down
Input
Control
Control
250
50
250
125
125
50
SR-3029
(nM)
eIF4G
eIF4E
β-actin

### Slide 4
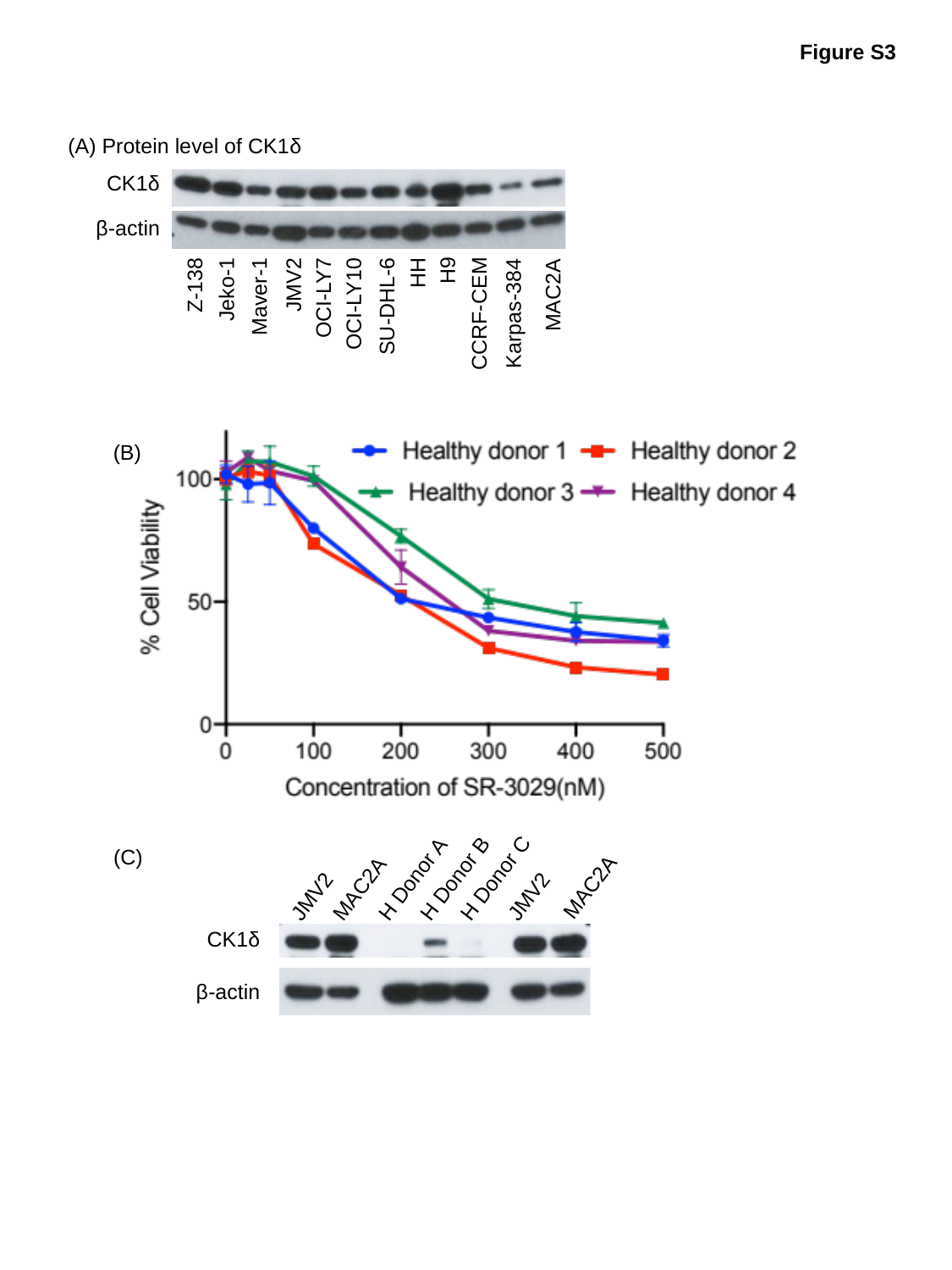

Figure S3
(A) Protein level of CK1δ
CK1δ
β-actin
HH
Z-138
H9
Jeko-1
Maver-1
OCI-LY7
JMV2
MAC2A
OCI-LY10
SU-DHL-6
CCRF-CEM
Karpas-384
(B)
(C)
MAC2A
MAC2A
H Donor C
H Donor B
JMV2
H Donor A
JMV2
CK1δ
β-actin

### Slide 5
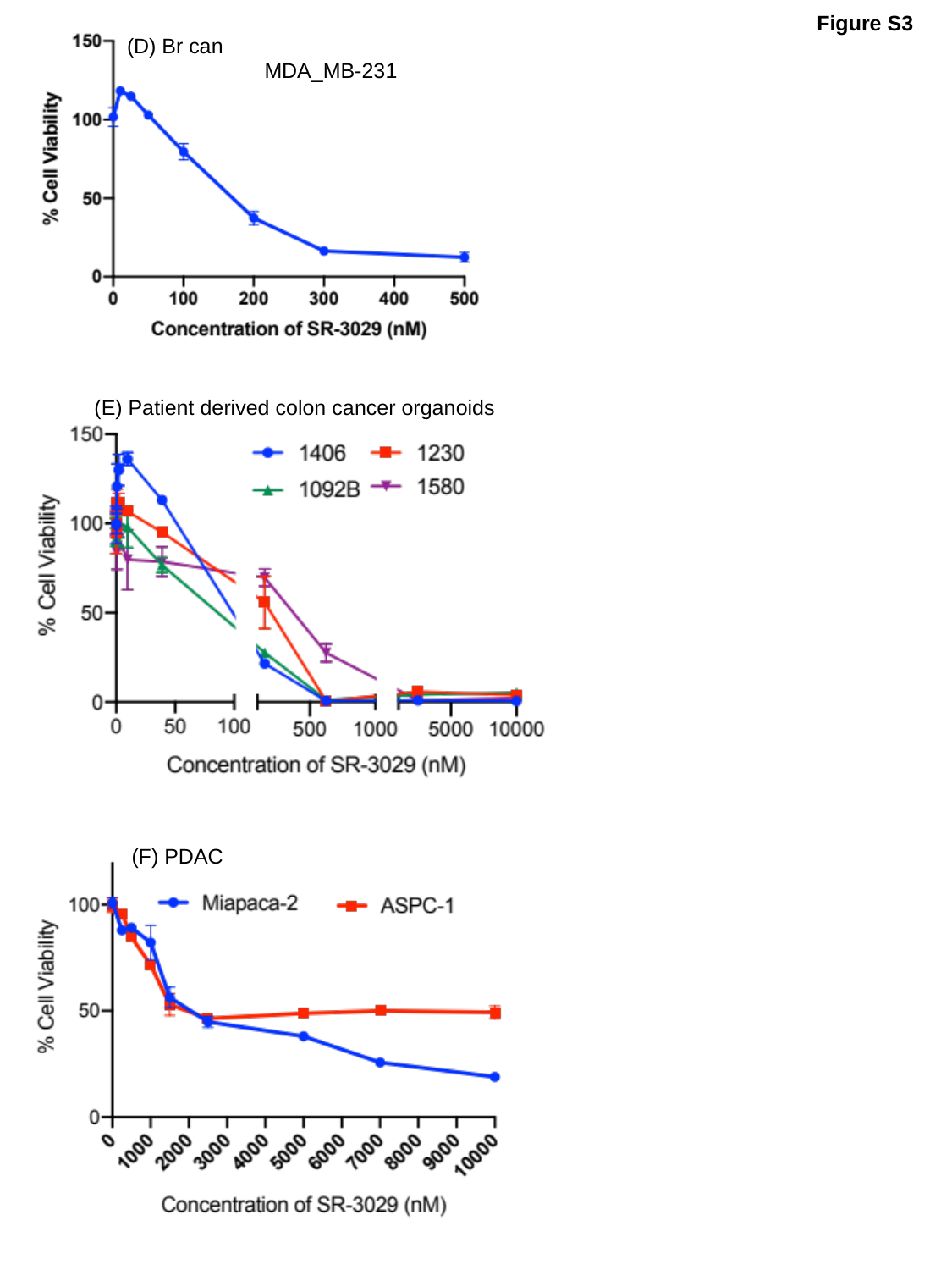

Figure S3
(D) Br can
MDA_MB-231
(E) Patient derived colon cancer organoids
(F) PDAC

### Slide 6
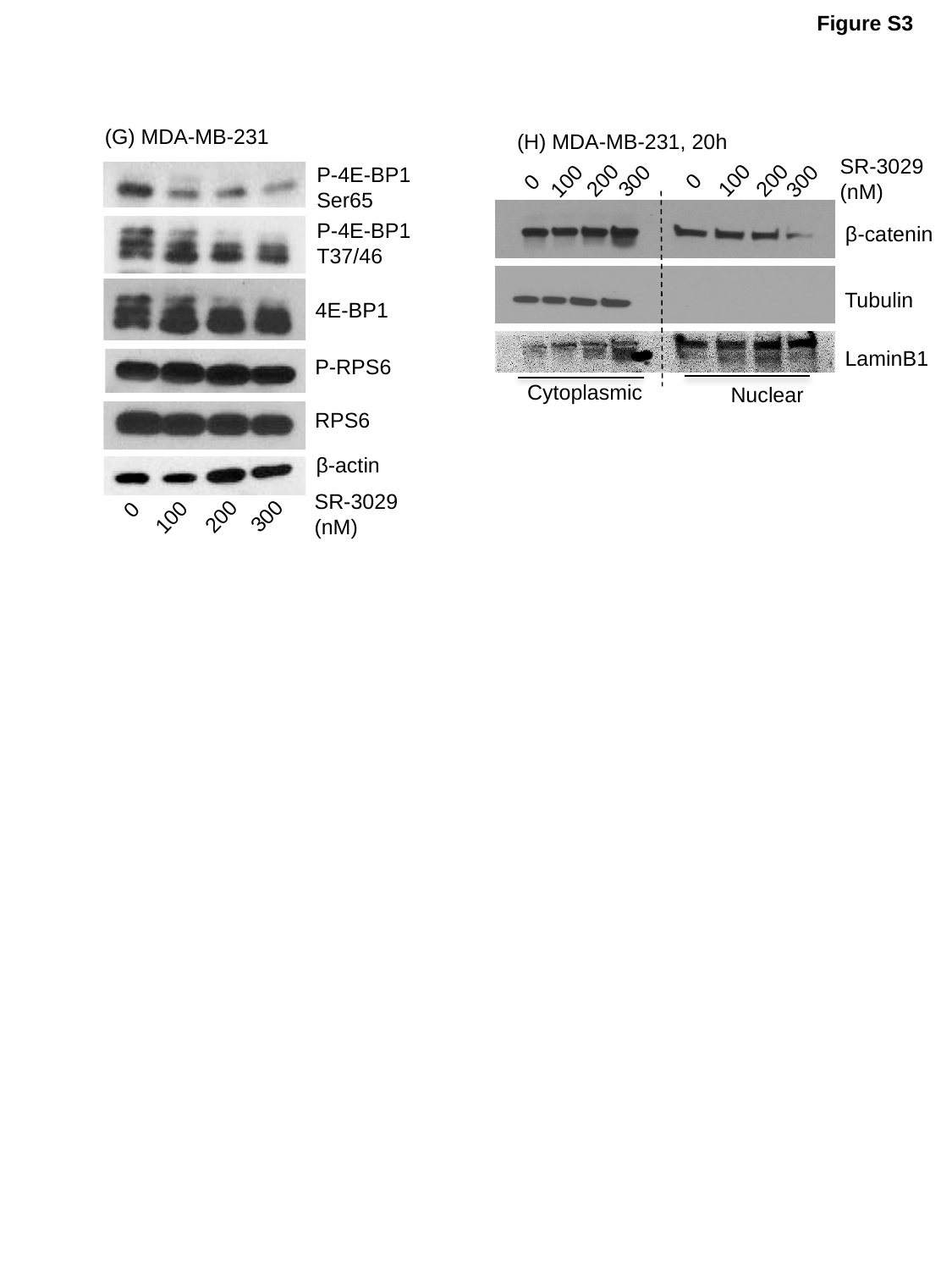

Figure S3
(G) MDA-MB-231
P-4E-BP1
Ser65
P-4E-BP1
T37/46
4E-BP1
P-RPS6
RPS6
β-actin
SR-3029
(nM)
0
300
200
100
(H) MDA-MB-231, 20h
SR-3029
(nM)
200
100
200
300
300
100
0
0
β-catenin
Tubulin
LaminB1
Cytoplasmic
Nuclear

### Slide 7
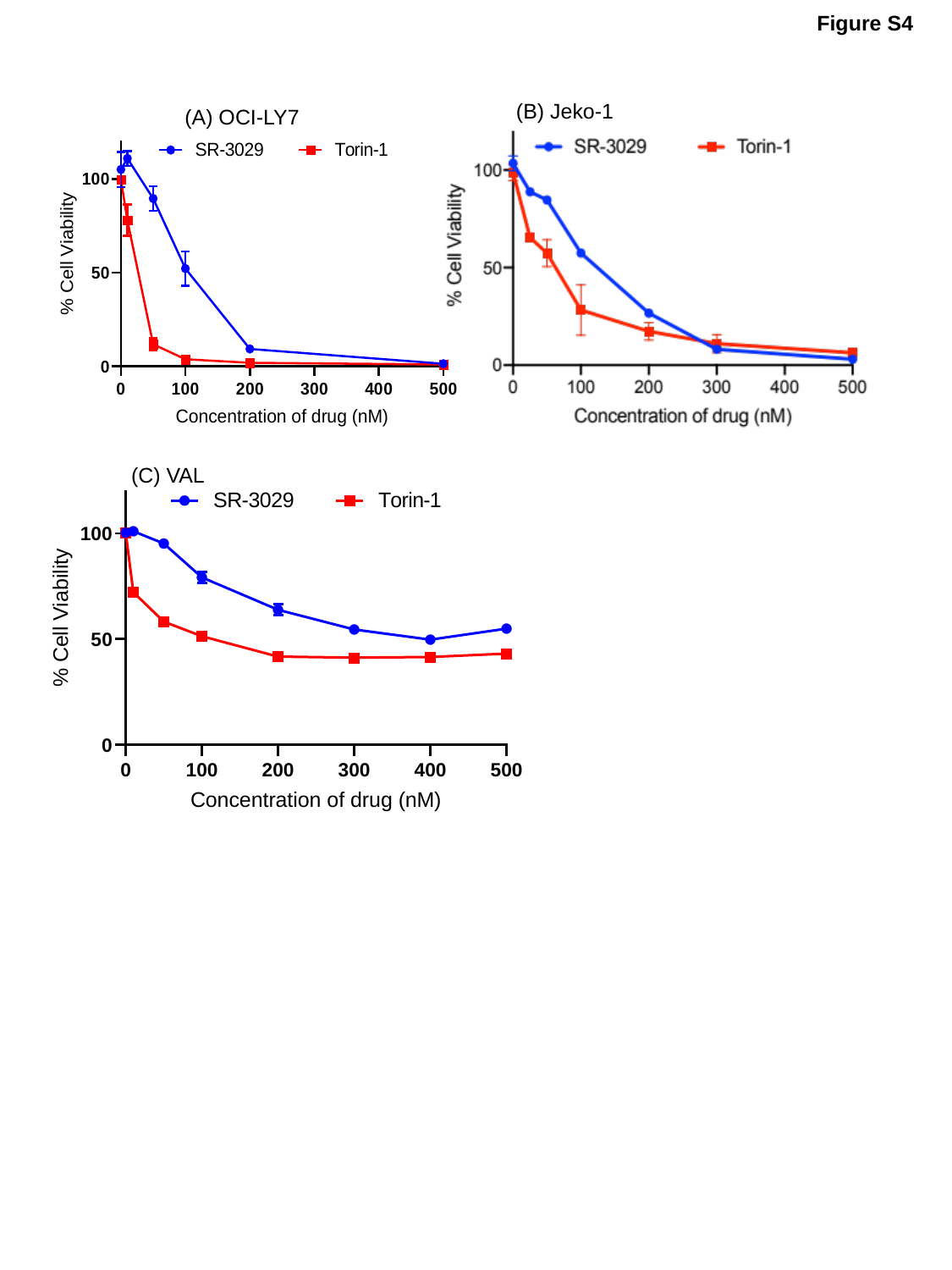

Figure S4
(B) Jeko-1
(A) OCI-LY7
(C) VAL

### Slide 8
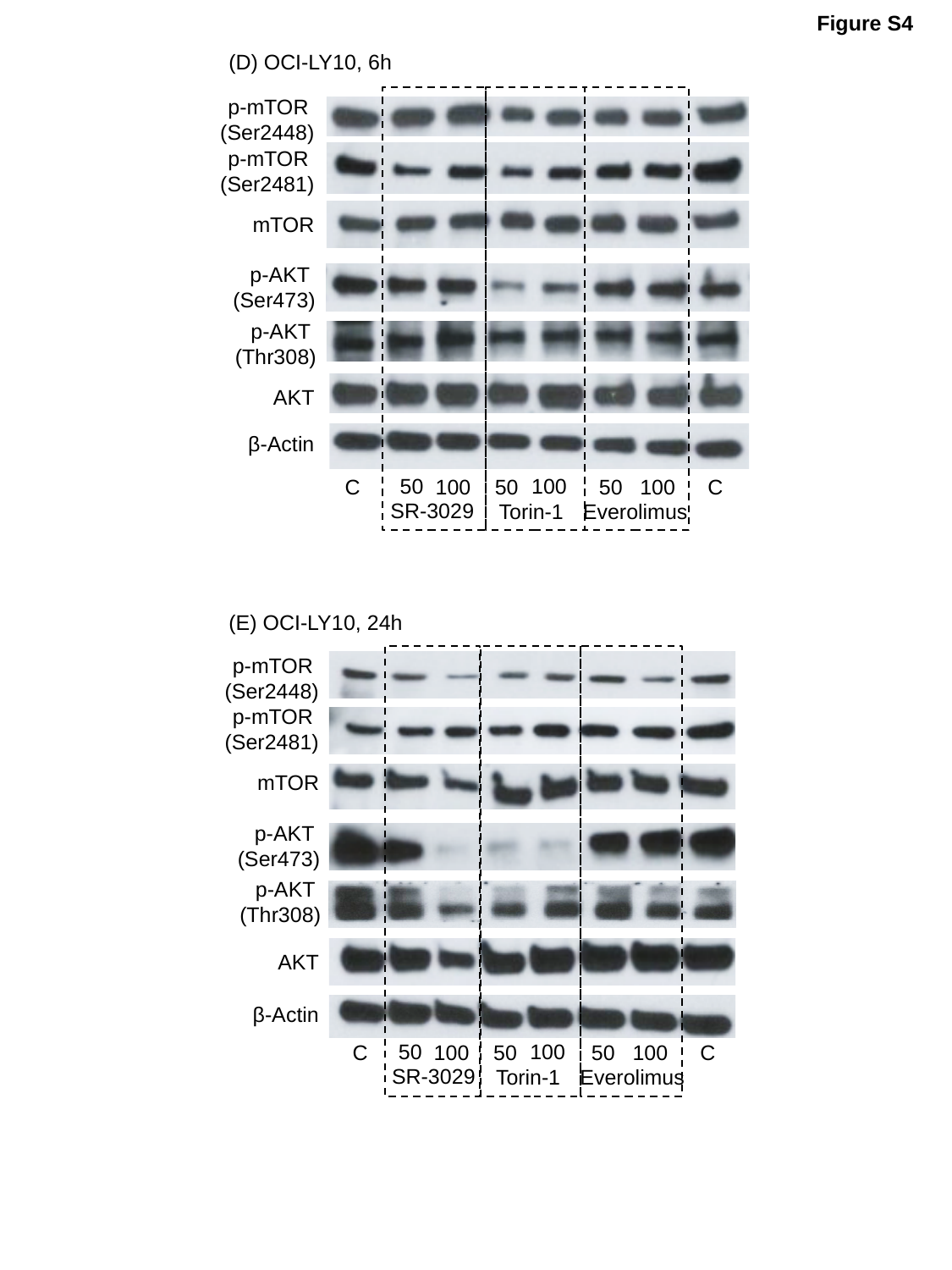

Figure S4
(D) OCI-LY10, 6h
p-mTOR
(Ser2448)
p-mTOR
(Ser2481)
mTOR
p-AKT
(Ser473)
p-AKT
(Thr308)
AKT
β-Actin
 100
50
 100
50
 C
 100
50
 C
SR-3029
Torin-1
Everolimus
(E) OCI-LY10, 24h
p-mTOR
(Ser2448)
p-mTOR
(Ser2481)
mTOR
p-AKT
(Ser473)
p-AKT
(Thr308)
AKT
β-Actin
 100
50
 100
50
 C
 100
50
 C
SR-3029
Torin-1
Everolimus

### Slide 9
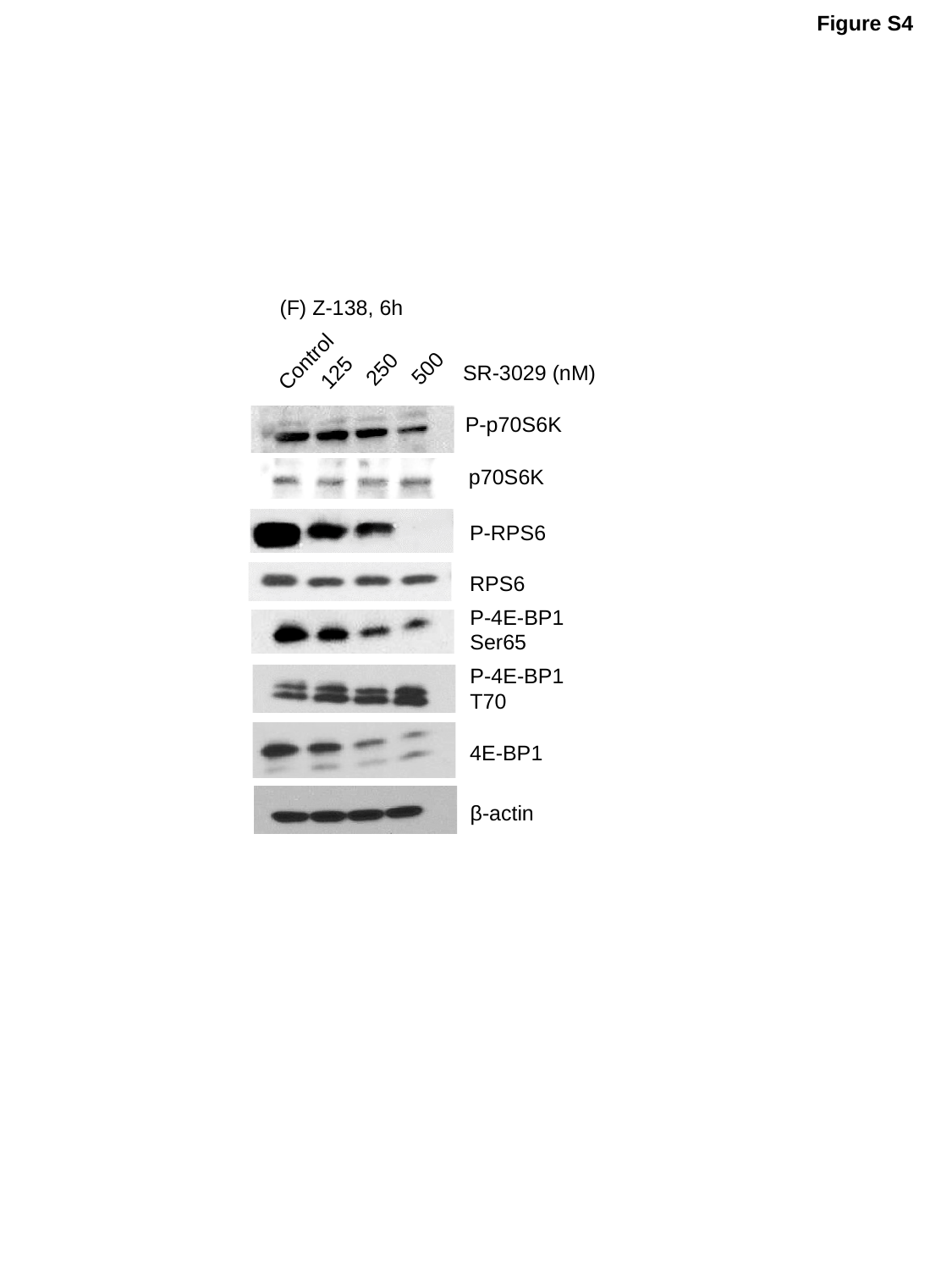

Figure S4
(F) Z-138, 6h
Control
500
250
125
SR-3029 (nM)
P-p70S6K
p70S6K
P-RPS6
RPS6
P-4E-BP1
Ser65
P-4E-BP1
T70
4E-BP1
β-actin

### Slide 10
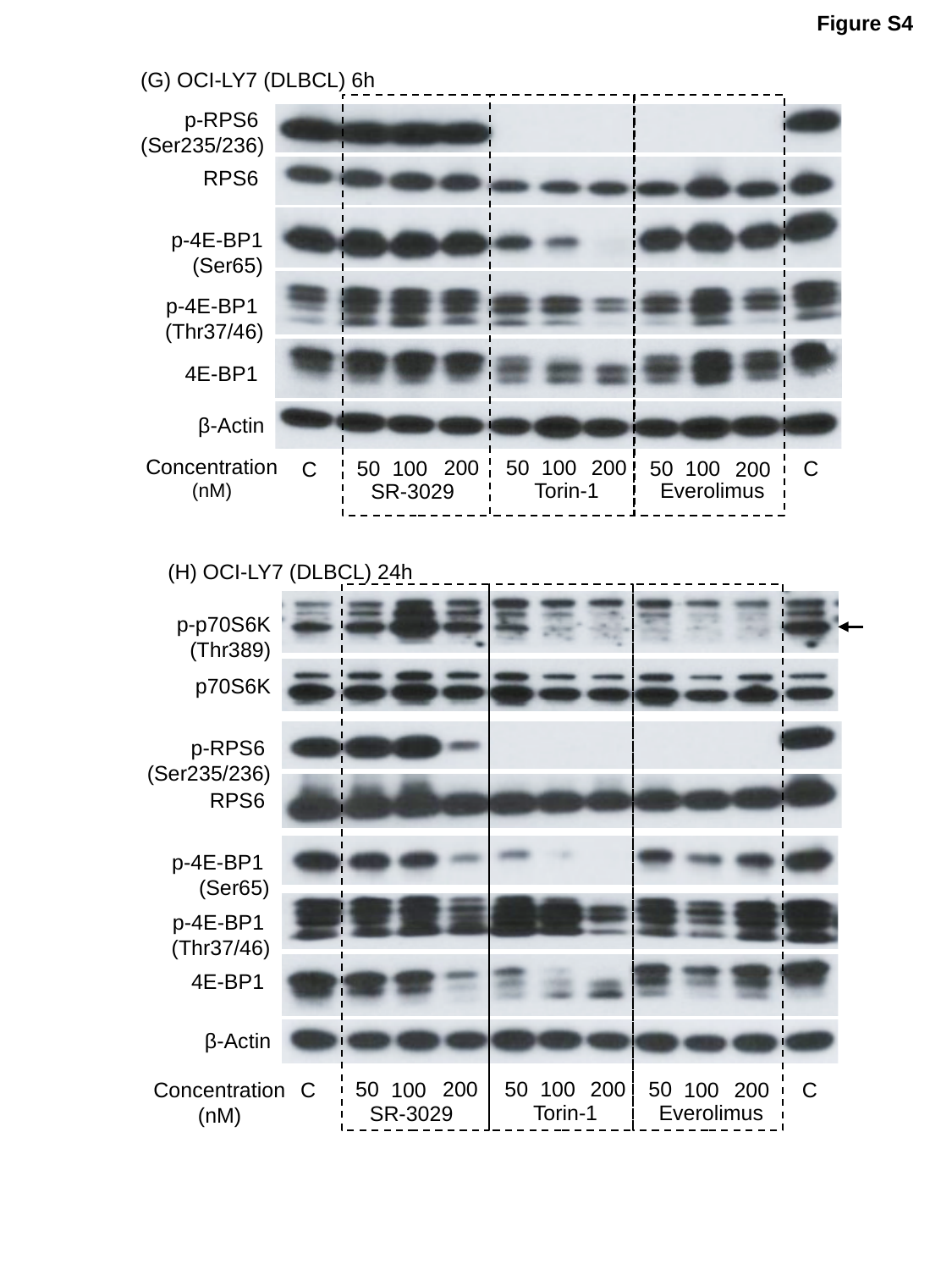

Figure S4
(G) OCI-LY7 (DLBCL) 6h
p-RPS6
(Ser235/236)
RPS6
p-4E-BP1
(Ser65)
p-4E-BP1
(Thr37/46)
4E-BP1
β-Actin
Concentration
(nM)
50
 200
 100
 200
50
50
 C
 100
 100
 C
 200
Torin-1
Everolimus
SR-3029
(H) OCI-LY7 (DLBCL) 24h
p-p70S6K
(Thr389)
p70S6K
p-RPS6
(Ser235/236)
RPS6
p-4E-BP1
(Ser65)
p-4E-BP1
(Thr37/46)
4E-BP1
β-Actin
50
 200
 100
 200
50
50
 C
 100
 100
Concentration
(nM)
 C
 200
Torin-1
Everolimus
SR-3029

### Slide 11
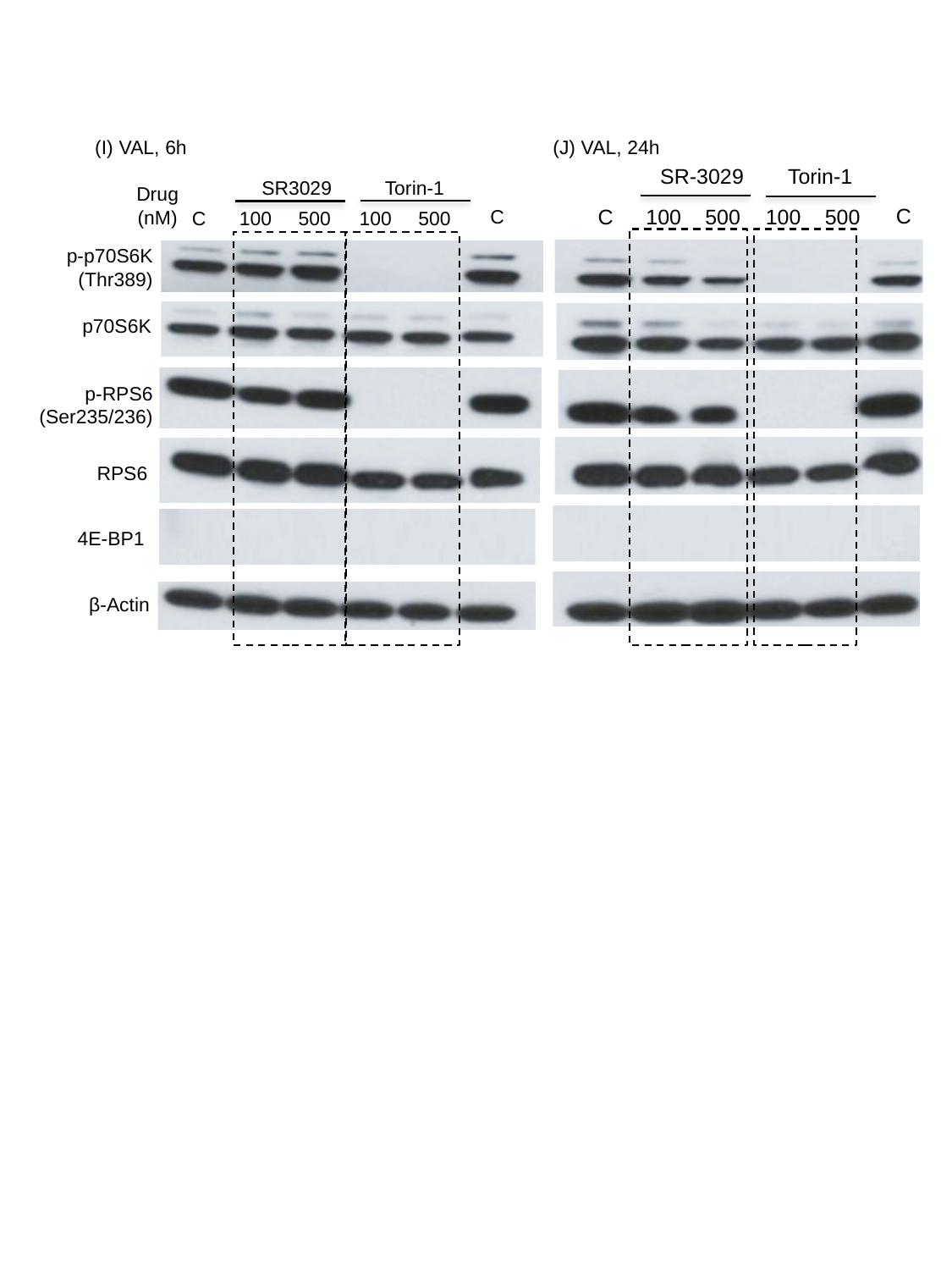

(J) VAL, 24h
SR-3029
Torin-1
 C
 500
 500
 100
 C
 100
(I) VAL, 6h
SR3029
Torin-1
Drug
(nM)
 C
 500
 500
 100
 C
 100
p-p70S6K (Thr389)
p70S6K
p-RPS6 (Ser235/236)
RPS6
4E-BP1
β-Actin

### Slide 12
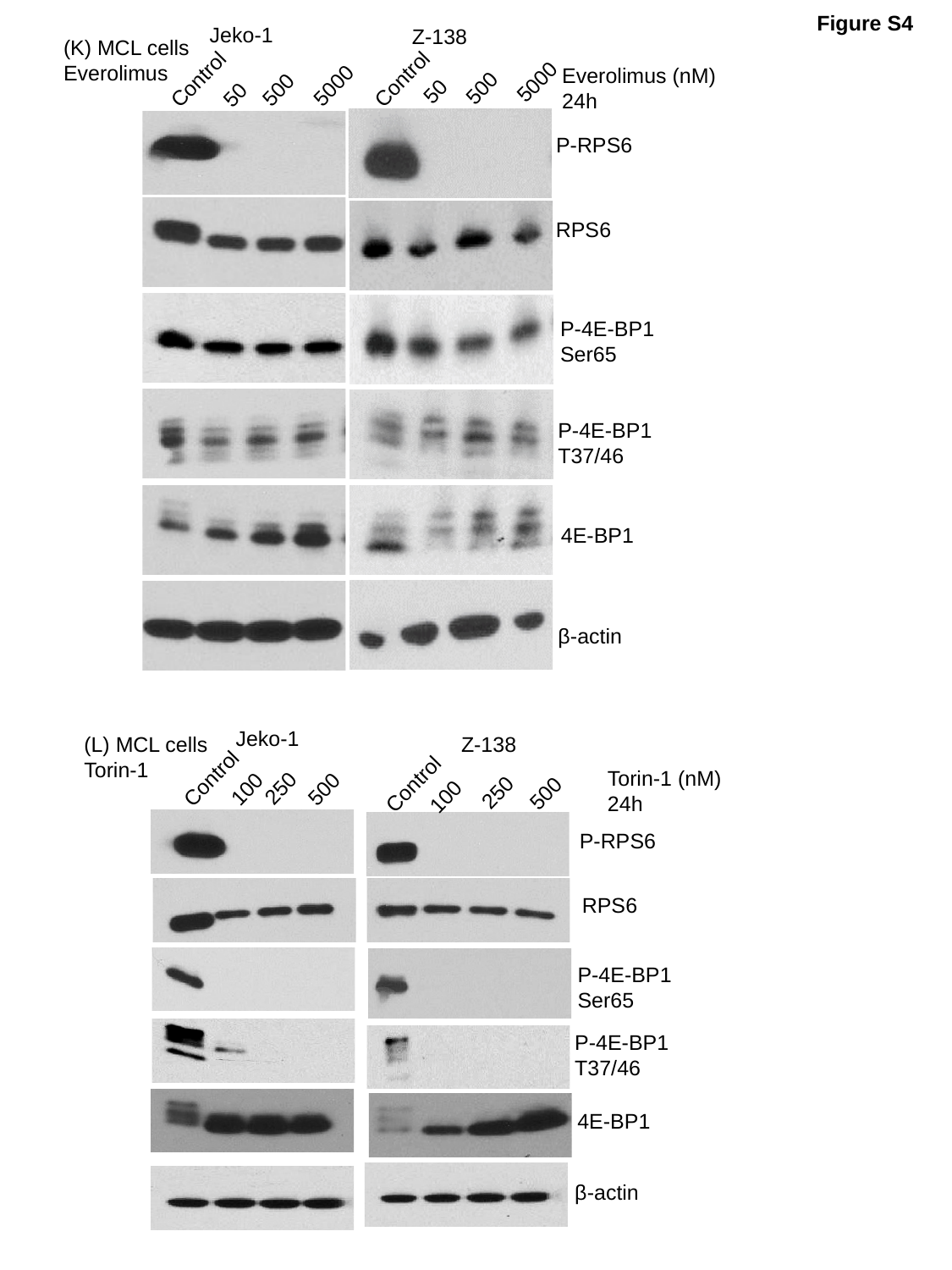

Figure S4
Jeko-1
Z-138
Everolimus (nM)
24h
Control
Control
5000
5000
500
500
50
50
P-RPS6
RPS6
P-4E-BP1
Ser65
P-4E-BP1
T37/46
4E-BP1
β-actin
(K) MCL cells
Everolimus
Jeko-1
Z-138
Torin-1 (nM)
24h
Control
Control
250
100
500
250
500
100
P-RPS6
RPS6
P-4E-BP1
Ser65
P-4E-BP1
T37/46
4E-BP1
β-actin
(L) MCL cells
Torin-1

### Slide 13
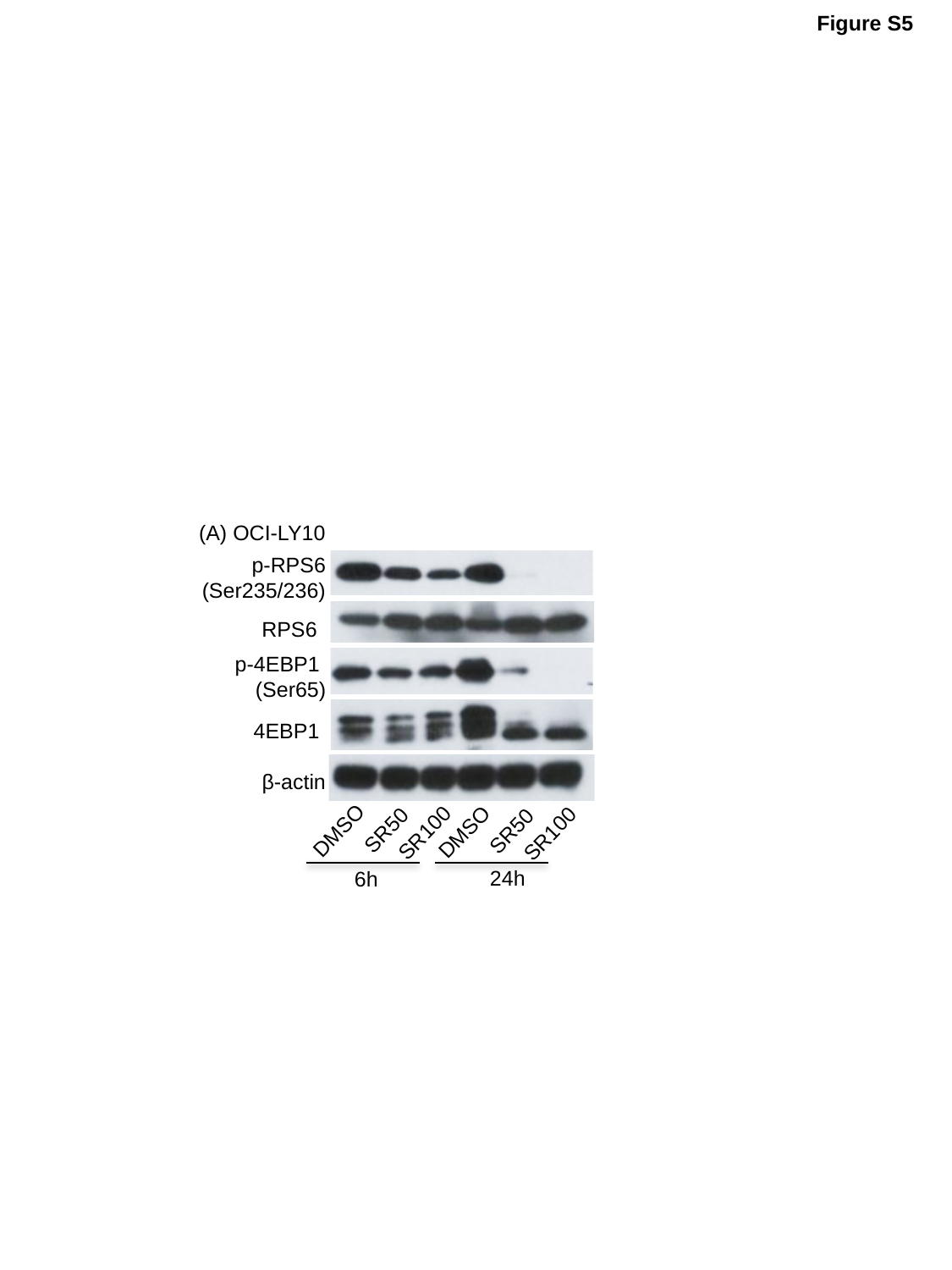

Figure S5
(A) OCI-LY10
p-RPS6 (Ser235/236)
RPS6
p-4EBP1
(Ser65)
4EBP1
β-actin
SR50
SR50
DMSO
DMSO
SR100
SR100
24h
6h

### Slide 14
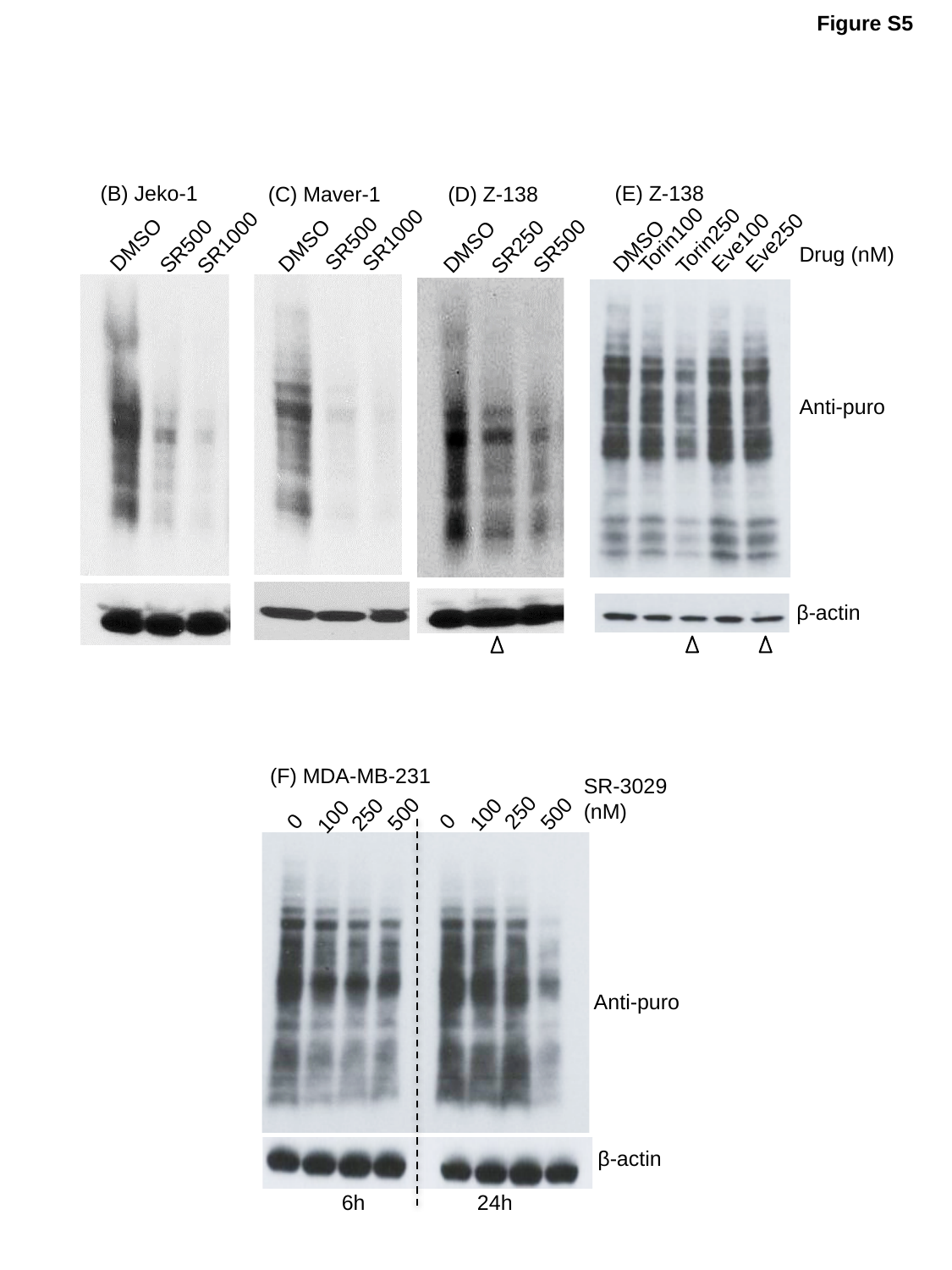

Figure S5
Torin100
Torin250
Eve250
Eve100
DMSO
(E) Z-138
(B) Jeko-1
SR1000
DMSO
SR500
(C) Maver-1
SR1000
SR500
DMSO
(D) Z-138
SR250
DMSO
SR500
Drug (nM)
Anti-puro
β-actin
(F) MDA-MB-231
SR-3029
(nM)
250
100
250
500
100
500
0
0
Anti-puro
β-actin
6h
24h

### Slide 15
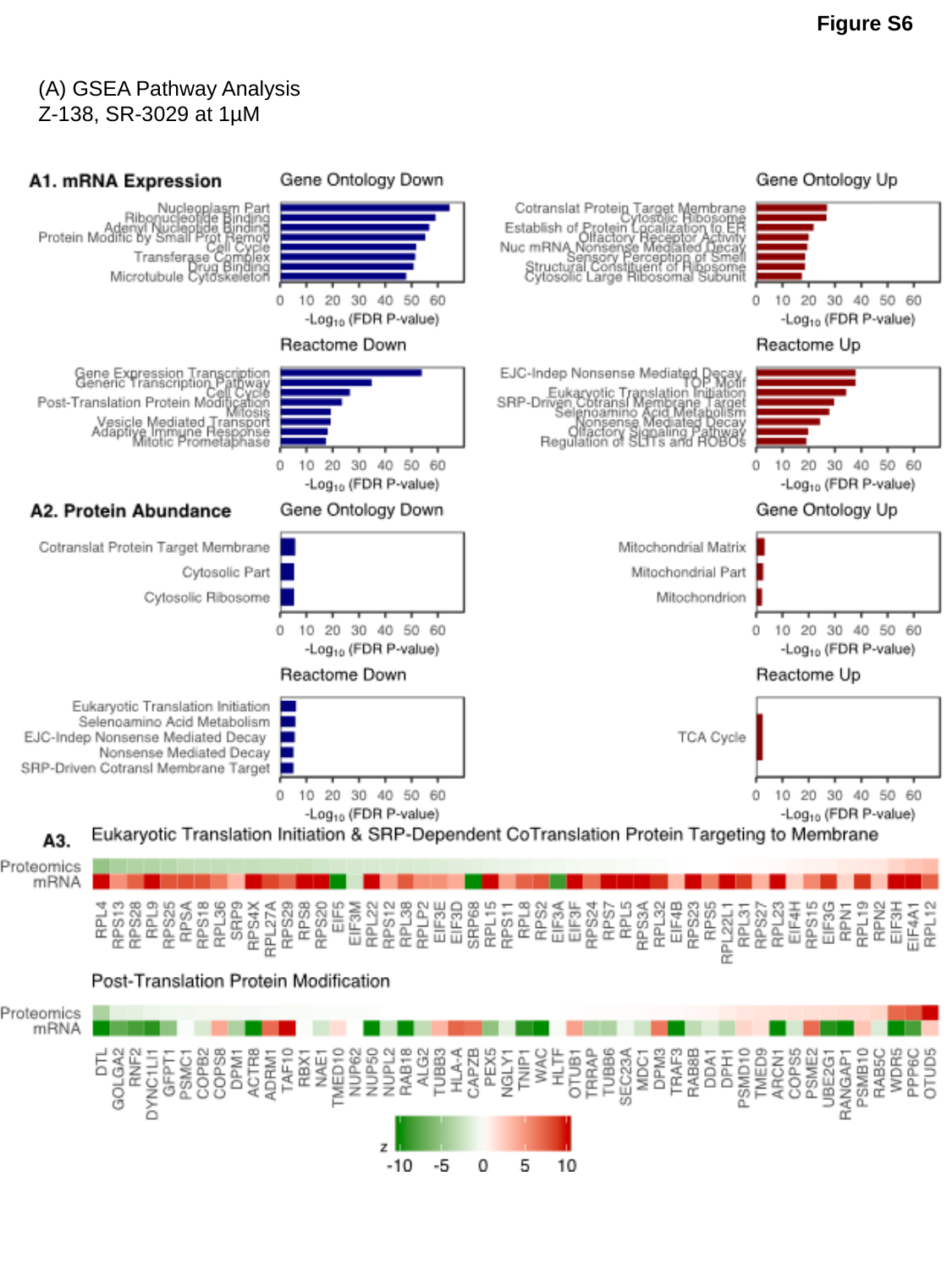

Figure S6
(A) GSEA Pathway Analysis
Z-138, SR-3029 at 1µM

### Slide 16
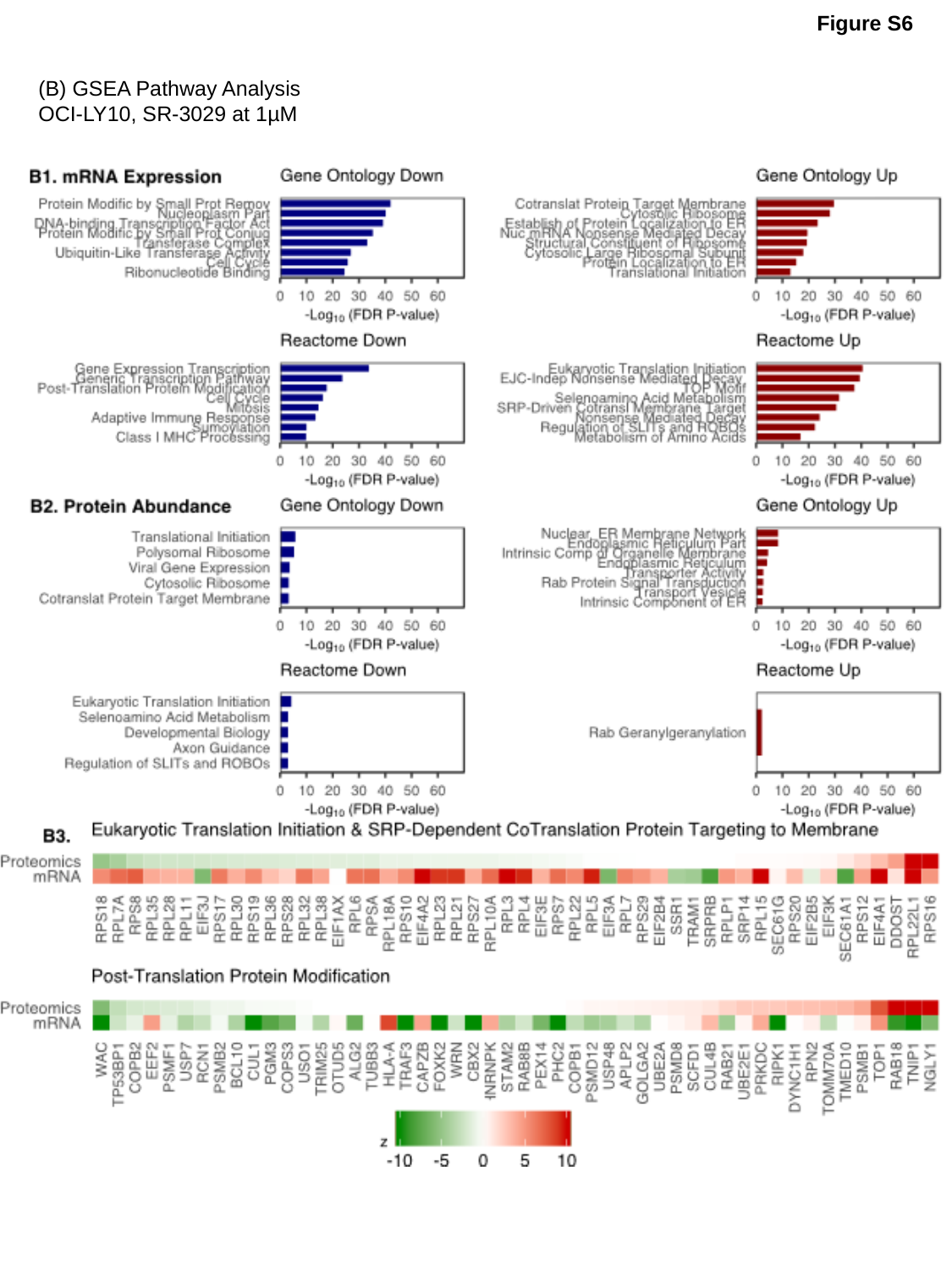

Figure S6
(B) GSEA Pathway Analysis
OCI-LY10, SR-3029 at 1µM

### Slide 17
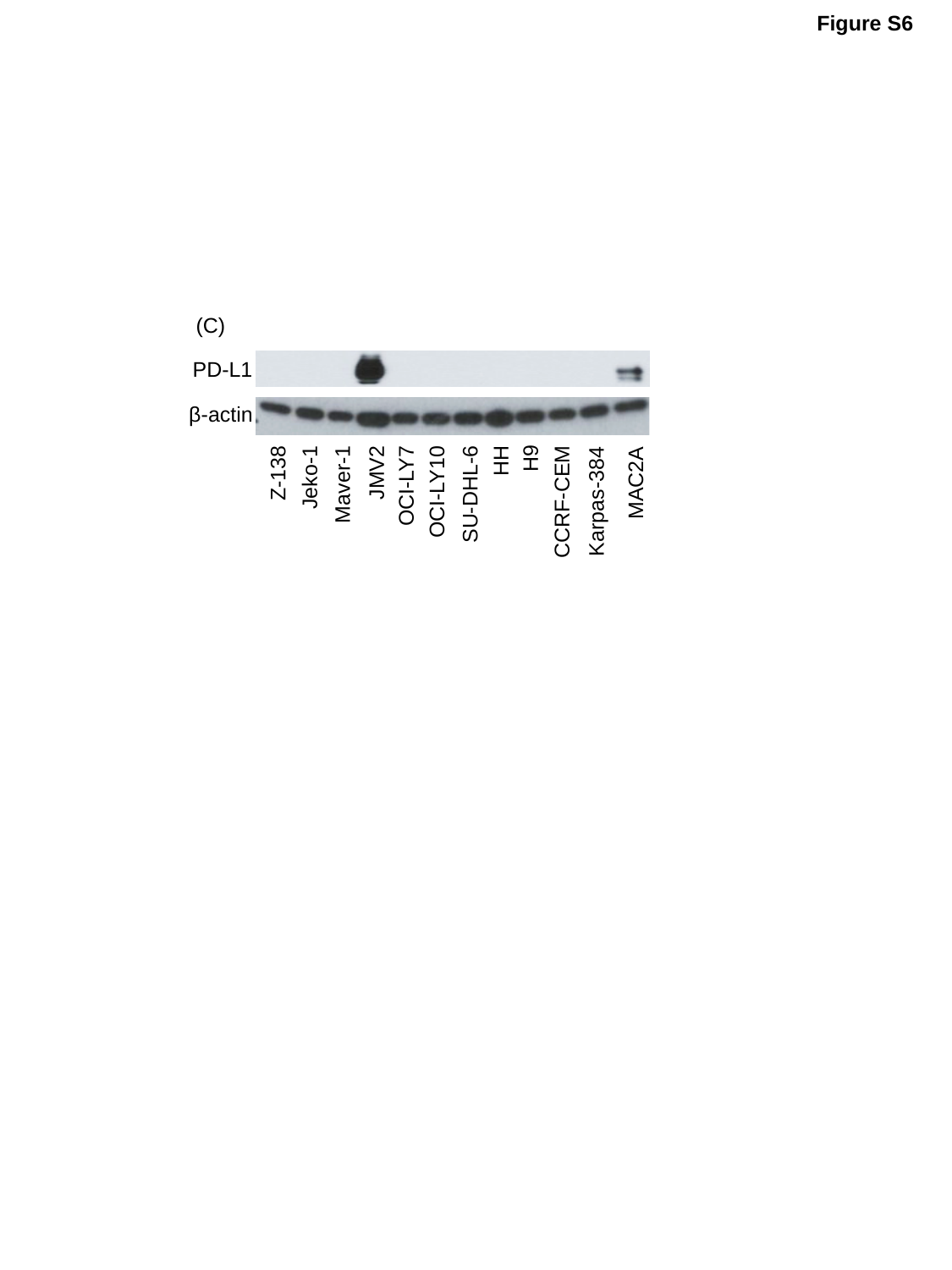

Figure S6
(C)
PD-L1
β-actin
HH
Z-138
H9
Jeko-1
Maver-1
OCI-LY7
JMV2
MAC2A
OCI-LY10
SU-DHL-6
CCRF-CEM
Karpas-384
