## Supplementary Files for "CK1 Delta Is an mRNA Cap-Associated Protein That Drives Translation Initiation and Tumor Growth"

**Supplemental figure legends**

**Table S1.** IC50 and IC90 values of SR-3029 at 24 and 48 in cancer and non-cancer cells, as determined by the Cell Titer Glo assay.

**Figure S1.** Knockout cell lines and confirmation. **(A)** Sequencing analysis of CK1δ-/- knockout clones in HEK293. **(B)** OCI-LY7 was targeted for CK1ε knockout by CRISPR-Cas9 system. The CK1ε knockout cells in LY7 were a mixed population. The cells were subjected to immunoblotting with the indicated antibodies.

**Figure S2.** Cap binding assay. **(A)** Jeko-1 cell line was treated with DMSO, SR-3029 50-250 nM for 24h then processed for the cap binding assay using m^7^GTP agarose beads to pull down cap binding proteins, which were then analyzed by immunoblotting. The ratio of eIF4G in the pulldown relative to the input cell lysate is an index of eIF4F assembly.

**Figu**r**e S3.** Expression of CK1δ and responses to SR-3029. **(A)** Protein expression profile of CK1δ in the indicated lymphoma cell lines, as determined by immunoblot. **(B)** Response of peripheral blood mononuclear cells (PBMCs) from healthy donors #1-4 to SR-3029 treatment for 48h, as determined by the Cell Titer Glo assay. **(C)** Protein level of CK1δ in PBMCs relative to lymphoma cell lines as determined by immunoblot. **(D)** Breast cancer cell line MDA-MB-231 was treated as indicated for 48h and analyzed using the Cell-Titer Glo assay. **(E)** Colon cancer organoids. Colon cancer cells from patients were plated as 2000 cells per well in a 96 well plate in 10 µL of Matrigel. After polymerization, 50 µL of organoid media was added to each well. Cells were allowed to grow for 3 days followed by drug treatment for 4 days. SR-3029 was added in a 3- or 4-point serial dilution. The viable cells in the treated and control samples were quantitated using the Cell-Titer Glo assay. **(F)** Two human cell lines of pancreatic ductal adenocarcinoma (PDAC) were treated as indicated for 24, 48, and 72h. Cell viability was calculated as above using the Cell-Titer Glo assay. The viability curves showed only the results from 72h. (**G-H**) Breast cancer cell line MDA-MB-231 was treated with DMSO or SR-3029 for 24h and processed for immunoblotting (G), or for 20h then processed for nuclear and cytoplasmic fractions and probed for the level of β-catenin (H). Tubulin and Lamin B1 were the controls for the cytoplasmic and nuclear proteins, respectively.

**Figure S4.** The cytotoxic and molecular effects of SR-3029 relative to mTOR inhibitors. **(A-C)** Cell viability curve of the OCI-LY7 (A), Jeko-1 (B), and VAL (C) cell lines were treated with SR-3029 or Torin-1 for 48h. Viable cells were determined by Cell Titer Glo assay, and the viability of the treated cells was expressed as the percentage compared to the vehicle control. **(D-J)** Various lymphoma cell lines were treated with SR-3029, Torin-1, Everolimus, or vehicle control at the indicated concentrations for 6h or 24h, then processed for immunoblotting. Arrow indicates the specific band of phospho-p70S6K (H). **(K-L)** Lymphoma cell lines Jeko-1 and Z-138 were treated with Everolimus (K) or Torin-1 (L), or DMSO for 24h and processed for immunoblotting.

**Figure S5.** Effects of SR-3029 on translation and translation regulators. **(A)** OCI-LY10 cells were treated with DMSO or SR-3029 (SR) at the indicated concentrations or the DMSO control for 6h or 24h. **(B-F)** Lympoma cell lines (B-E) and breast cancer cell line MDA-MB-231 (F) were treated with SR-3029 (SR), Torin, Everolimus (Eve) at the indicated concentrations or the DMSO control for 6h or 24h. Puromycin was then added at the final concentration of 1μg/ml for 30 min. Cell lysates were subjected to Western blotting using the anti-puromycin or anti-β-actin antibody. The empty triangles point to samples treated with 250 nM of the indicated drugs.

**Figure S6.** Effects of SR-3029 on gene expression genome wide and specific cancer related genes. **(A)** Pathway Analysis of the Effects of SR-3029 on gene and protein expression in Z-138. (A1-A2) Z-138 lymphoma cells were treated with 1 µM SR-3029 or DMSO control for 3h, and processed for RNAseq and proteomics studies. The RNA and protein abundance of each gene were compared between the treatment and control samples. Pathway analysis was performed on the SR-3029 induced differential RNA expression signature (A1) and the proteomics expression signature (A2) using the analytic rank-based enrichment analysis algorithm (a version of preranked GSEA). The most enriched Gene Ontology pathways and Reactome pathways are shown, ranked by FDR-corrected p-value, with down and up pathways displayed separately. (A3) Upper panel: Heatmap of the top 50 most differentially expressed genes/proteins involved in the Reactome Eukaryotic Translation Initiation pathway [R-HSA-72613] and SRP-dependent cotranslational targeting of proteins to ER/cell membrane [R-HSA-1799339]. As was seen at the 3 uM dose (main Figure 6), SR-3029 decreased the abundance of 40S ribosomal proteins but increased the mRNA expression of these genes. Lower panel: Heatmap of the top 50 most differentially expressed genes/proteins in the Reactome post-translational modification pathway [R-HSA-597592]. **(B)** Pathway Analysis of the Effects of SR-3029 on gene and protein expression in OCI-LY10. (B1-B2) OCI-LY10 lymphoma cells were treated with 1 µM SR-3029 or DMSO control for 3h, and processed for RNAseq and proteomics studies. The RNA and protein abundance of each gene were compared between the treatment and control samples. Pathway analysis was performed on the SR-3029 induced differential RNA expression signature (B1) and the proteomics expression signature (B2) using the analytic rank-based enrichment analysis algorithm (a version of preranked GSEA). The most enriched Gene Ontology pathways and Reactome pathways are shown, ranked by FDR-corrected p-value, with down and up pathways displayed separately. (B3) Upper panel: Heatmap of the top 50 most differentially expressed genes/proteins involved in the Reactome Eukaryotic Translation Initiation pathway [R-HSA-72613] and SRP-dependent cotranslational targeting of proteins to ER/cell membrane [R-HSA-1799339]. As was seen in Z-138 cells (main Figure 6), SR-3029 decreased the abundance of 40S ribosomal proteins but increased the mRNA expression of these genes. Lower panel: Heatmap of the top 50 most differentially expressed genes/proteins in the Reactome post-translational modification pathway [R-HSA-597592]. **(C)** Protein expression profile of PD-L1 in the indicated lymphoma cell lines.
